# Chemical Systems Biology Reveals Mechanisms of Glucocorticoid Receptor Signaling

**DOI:** 10.1101/2020.06.15.153270

**Authors:** Nelson E. Bruno, Jerome C. Nwachukwu, Sathish Srinivasan, Charles C. Nettles, Tina Izard, Zhuang Jin, Siddaraju V. Boregowda, Donald G. Phinney, Olivier Elemento, Xu Liu, Eric A. Ortlund, René Houtman, Diana A. Stavreva, Gordon L. Hager, Theodore M. Kamenecka, Douglas J. Kojetin, Kendall W. Nettles

**Affiliations:** Department of Integrative Structural and Computational Biology, The Scripps Research Institute, Jupiter, FL 33458 USA; Department of Molecular Medicine, The Scripps Research Institute, Jupiter, FL 33458 USA; Caryl and Israel Englander Institute for Precision Medicine, Institute for Computational Biomedicine, Department of Physiology and Biophysics, Weill Cornell Medicine, New York, NY 10065, USA; Department of Biochemistry, Emory University School of Medicine, Atlanta, GA 30322, USA; Department of Research and Development, PamGene, 's-Hertogenbosch, 5211 HH, the Netherlands; Laboratory of Receptor Biology and Gene Expression, Building 41, B602, 41 Library Dr., National Cancer Institute, NIH, Bethesda, MD 20892-5055, USA

## Abstract

Glucocorticoids display remarkable anti-inflammatory activity, but their use is limited by on-target adverse effects including insulin resistance and skeletal muscle atrophy. We used a chemical systems biology approach, Ligand Class Analysis (LCA), to examine ligands designed to modulate glucocorticoid receptor activity through distinct structural mechanisms. These ligands displayed diverse activity profiles, providing the variance required to identify target genes and coregulator interactions that were highly predictive of their effects on myocyte glucose disposal and protein balance. Their anti-inflammatory effects were linked to glucose disposal but not muscle atrophy. This approach also predicted selective modulation *in vivo*, identifying compounds that were muscle sparing or anabolic for protein balance and mitochondrial potential. LCA defined the mechanistic links between the ligand-receptor interface and ligand-driven physiological outcomes, a general approach that can be applied to any ligand-regulated allosteric signaling system.

## Introduction

Glucocorticoids (GCs) are among the most prescribed medicines due to their remarkable anti-inflammatory effects. Efforts to develop improved GCs have been hampered by poor understanding of the structural and molecular mechanisms through which glucocorticoid receptor (GR) ligands drive diverse phenotypic outcomes such as effects on inflammation, glucose disposal, or skeletomuscular atrophy^1,2^. This phenomenon is called selective modulation, and is a common feature of allosteric signaling through GR and other nuclear receptor superfamily transcription factors^3^, G protein-coupled receptors, and other small molecule drug targets.

The adrenal gland secretes GCs in a circadian fashion and in an acute response to nutrient deprivation or other stressors. In skeletal muscle, GCs stimulate catabolic signaling that opposes the insulin/PI3K/AKT signaling pathway, inhibits glucose uptake, and releases amino acids by inhibiting protein synthesis and stimulating protein degradation^2^. Upon GC binding, GR dissociates from inhibitory heat shock protein complexes, translocates to the nucleus, oligomerizes upon binding to DNA, and recruits an ensemble of coregulators, including chromatin-remodeling and histone-modifying enzymes that regulate gene expression. Several GR coregulators and target genes have been identified in skeletal muscle^4–7^, but factors that drive selective modulation are unclear.

We developed a chemical systems biology approach called Ligand Class Analysis (LCA) to identify molecular mechanisms that drive ligand-selective signaling^8–10^. With LCA, we characterize a series of related compounds that display a full range of variance in phenotypes such as insulin-mediated glucose uptake or protein synthesis and use that variance to define molecular predictors of the phenotype. This enables us to identify ligand-dependent gene expression and protein-interaction profiles that predict the molecular phenotypes of ligand classes that perturb receptor structure differently^8–10^. We then overexpress or knockdown predictive genes or interacting proteins to identify factors required for ligand-dependent phenotypes.

In this work, we evaluated the physiological effects of GCs during overnight nutrient deprivation when they coordinate acute metabolic adaptations, including glucose disposal and protein balance. We tested a GC series with different modifications on the steroidal scaffold^11^, providing distinct strategies to perturb GR structure and dynamics. Using a machine learning approach, we found that GCs selectively modulate protein balance in skeletal muscle versus glucose disposal and the inflammatory response. This identified coregulators and target genes that drive these activities, and enabled discovery of orally available GCs with muscle-sparing activity profiles in vivo.

## Results

### A GC-profiling platform for effects on skeletal muscle

We designed a series of GCs to test three different structural effects of PF-0251802 (PF802) (**1**), a selective modulator, the pro-drug of which reached clinical trials (**Extended Data Fig. 1a**)^12,13^. PF802 (**1**) drives antagonism by disrupting the dexamethasone (Dex) (**2**)-bound GR conformation (**Extended Data Fig. 1b**), using its benzyl group which is similar to the carbon-11 (C11) substitution in the antagonist, RU-486 (**3**) (**Extended Data Fig. 1c**). To increase affinity, PF802 (**1**) has a trifluoromethyl group attached at the steroid C17-equivalent position, while RU-486 (**3**) and momethasone furoate (**4**) have other substitutions at this site (**Extended Data Fig. 1c-d**). PF802 (**1**) also contains a methylpyridinyl acetamide group attached at the steroid C3-equivalent position. This group presumably enters the solvent channel underneath the coregulatory binding site, as first seen with deacylcortivazol (**5**) (**Extended Data Fig. 1e**), to alter the shape of AF2, and the ensemble of coregulators that bind this surface, thus driving selective modulation as we previously showed for the estrogen receptor^9,14^.

To understand how these different substitutions control GR activity, we generated 22 related compounds based on a steroidal core, which enabled us to direct substitutions within the ligand-binding pocket (**Extended Data Fig. 1f**). These included substitutions of the methylpyridinyl acetamide at C3 to perturb the AF2 surface (**6-12**)(**Fig. 1a**); substitutions at C11 to drive agonism/antagonism (**13-20**) (**Fig. 1b**); substitutions at C17 to optimize affinity (**21-24**) (**Fig. 1c**); and other modifications (**Fig. 1d**). We have published the synthesis and preliminary structure-activity relationships for these compounds^11^.

**Figure 1.**
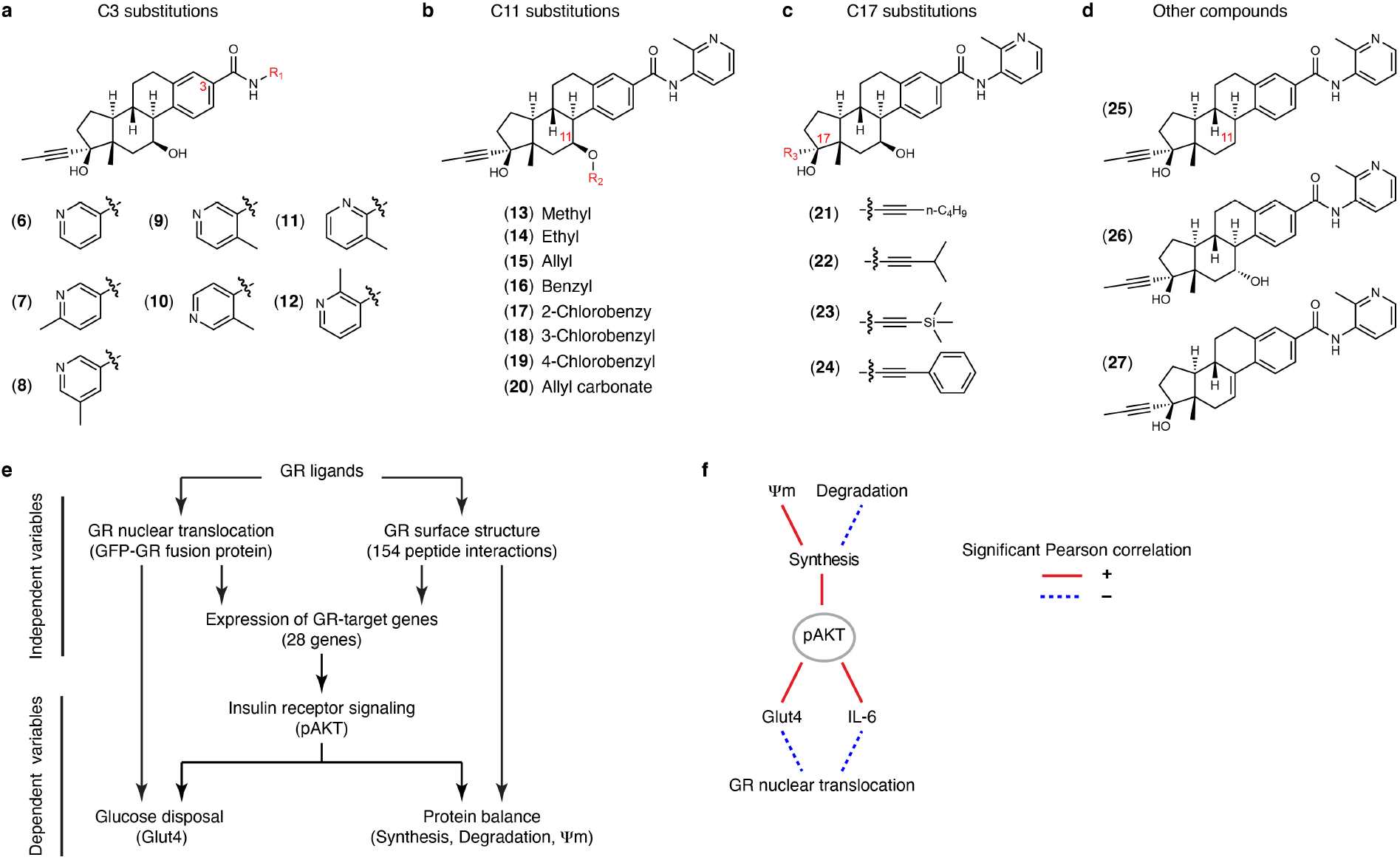
Structure-based design approach and glucocorticoid profiling platform. **a-d**) Glucocorticoids used in this study. **e)** Compound profiling and computational strategy. Effects on the independent variables were used in a machine learning approach, Random Forest, to identify predictors of skeletal muscle phenotypes, the dependent variables, described in **Figure 2**, **Supplementary Table 2,** and below. **f)** Relationships among GR-mediated phenotypes. Lines indicate significant Pearson correlation between variables using Bonferroni p_Adj_ < 0.0071.

We developed a compound profiling platform to examine how ligand-regulated GR nuclear localization, structure, and target gene expression in skeletal muscle control glucose disposal and protein balance (**Fig. 1e**), and whether these effects could be dissociated from anti-inflammatory effects such as the suppression of IL-6 secretion. We assayed ligand-induced nuclear translocation of a GFP-GR fusion protein, and probed for ligand-selective changes in surface structure by characterizing the GR-interaction profiles of 154 peptides derived from nuclear receptor coregulators and other interacting proteins using the MARCoNI FRET assay15. We also completed nascent RNA expression profiling of C2C12 myotubes, one hour after vehicle or Dex (**2**) treatment and selected 28 differentially expressed genes as potential regulators of glucose disposal and protein balance (see **Supplementary Information, Supplementary Table 1** for selection rationale). We compared ligand-dependent regulation of these genes in C2C12 myotubes using the nanoString nCounter for direct multiplexed transcript counting without PCR amplification, enabling highly quantitative and reproducible assays^16^. These datasets comprise the independent variables used in a machine learning approach to identify ligand-selective regulators of glucose disposal and protein balance in skeletal muscle, the dependent variables.

To assay physiologically relevant effects of GCs on skeletal muscle, myotubes were treated in the context of nutrient deprivation and a brief insulin challenge. One of the barriers to understanding GC action in skeletal muscle is that effects on protein balance required 50–100 μM Dex (**2**) *in vitro* ^17^, despite a ~5 nM Kd for GR. Effects on protein balance and Glut4 translocation also vary *in vivo*, depending upon the stressor. For example, GCs induce skeletal muscle atrophy and insulin resistance during fasting, while exercise has opposite effects^18,19^. We found that overnight serum-deprivation enabled robust GC-induced signal to noise after a brief insulin challenge with maximal effects of Dex (**2**) in the 1-10 nM range (**Methods, Extended Data Fig. 2a–g**). This suggests that GR acts by inhibiting the insulin/PI3K/AKT signaling pathway^20^.

We developed a compound profiling assay for phosphorylation of AKT at Thr308 (pAKT), due to its joint role in coordinating protein synthesis and glucose disposal (**Extended Data Fig. 2a–b**). For effects of compounds on glucose disposal, we measured the rate limiting step—insulin-mediated translocation of the glucose transporter, Glut4, to the cell surface. (**Extended Data Fig. 2c**). For protein balance, we developed two assays: 1) Degradation: GC-induced protein degradation assayed by the release of tritiated phenylalanine (**Extended Data Fig. 2d**); and 2) Synthesis: effects of the ligands on insulin-stimulated protein synthesis assayed using the SUnSET assay, which measures the incorporation of puromycin into nascent polypeptide chains (**Extended Data Fig. 2e**). We assessed the relationship between protein balance and mitochondrial potential (Ψm) measured by high content imaging after staining with a MitoTracker dye, where accumulation of the dye is dependent on membrane potential, and modulated by the GR ligands. (**Extended Data Fig. 2f–h**). We also compared ligand-dependent effects on IL-1β-induced secretion of IL-6 by A549 lung cells, a standard assay for the inflammatory response. Ligand activity in these assays were used as dependent variables to identify predictive coregulator interactions and target genes, and selective modulators.

We identified two clusters of dependent variables, connected by the effects of the ligands on pAKT. In the first cluster, effects of the ligands were significantly intercorrelated between effects on Synthesis, Degradation, pAKT, and ψm, suggesting transcriptional control via a common set of coregulators and target genes (**Fig. 1f**, **Extended Data Fig. 2i**, **Supplementary Dataset** shows the Pearson correlation matrix and P values). In the second cluster, GC effects on IL-6 and Glut4 were correlated to pAKT and GR nuclear translocation (**Fig. 1f**). However, GC effects on pAKT and GR nuclear translocation were not correlated (**Extended Data Fig. 2j**), indicating that some ligand-dependent outcomes of GR signaling (e.g. Glut4, IL-6) involve a graded response to the amount of nuclear GR, while others require a threshold amount of nuclear GR. These results suggest that GCs modulate different transcriptional signaling pathways— coregulators and target genes—that coordinate some processes but allow others to be dissociated.

### Machine learning defines top predictors of GC action

We performed predictive modeling of each GR-mediated phenotype (dependent variable) using *Random Forest*, a classification and regression tree algorithm^21^ (**Fig. 1e**). To identify a statistically significant set of gene expression and peptide interaction predictors for each phenotype, we used *Boruta*, a Random Forest-based wrapper algorithm^22^. We iteratively compared the importance of each feature (gene or peptide) in the dataset to a “shadow” dataset obtained by shuffling the dependent and independent variables. In each of 500 iterative decision rounds, a feature was selected if it was more significant than all the shadow features. We then identified a minimal set of predictors by applying a forward selection strategy. The set of significant features were tested one-by-one, starting with the most significant, for their ability to improve the model. Features that improved the model were retained, while those that worsened the model were removed. With this approach, we were able to predict between 47% (Degradation) and 84% (Glut4) of the variance in the dependent variables (**Fig. 2a**, **Supplementary Table 2**, **Supplementary Dataset**).

**Figure 2.**
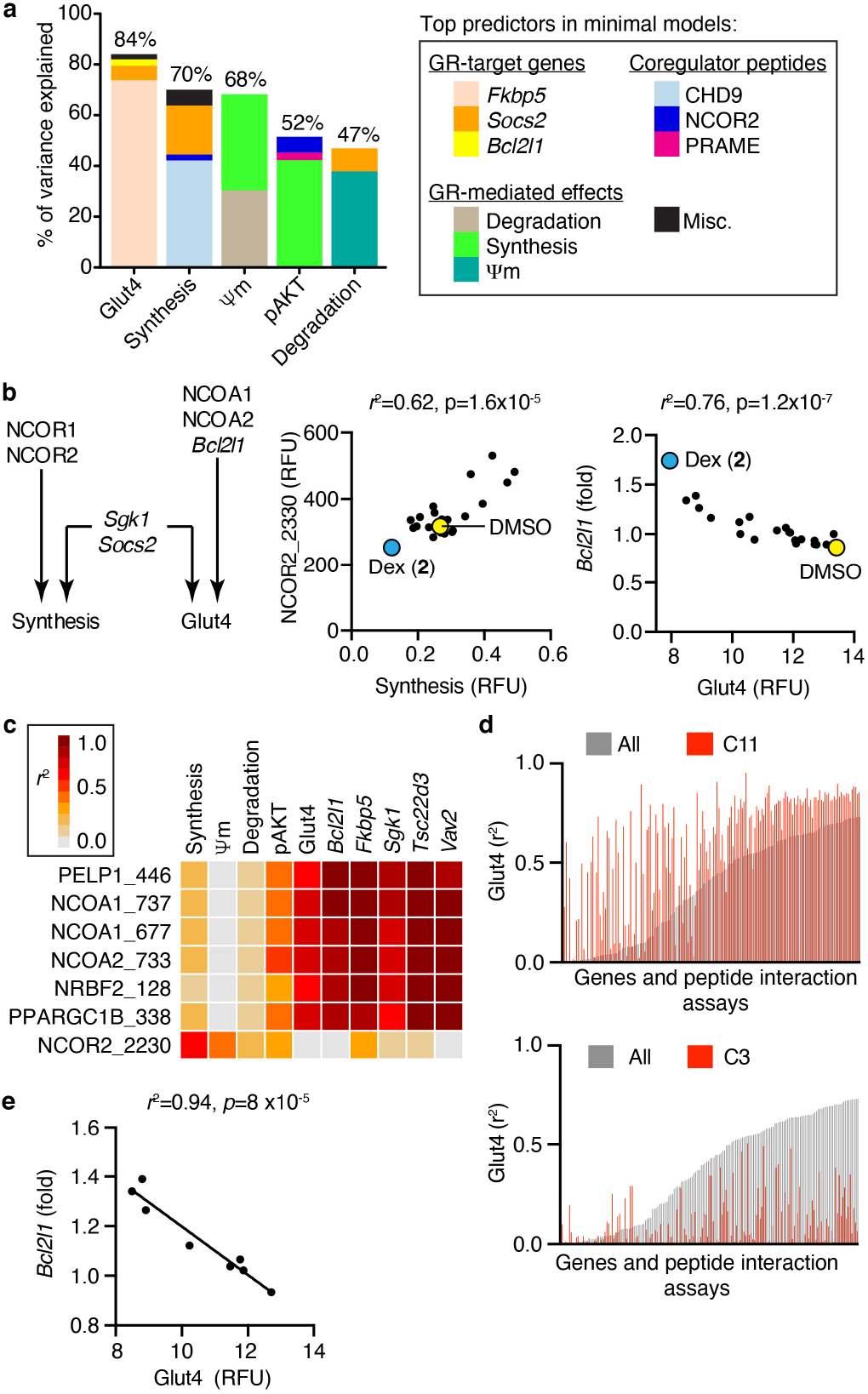
Machine learning reveals top predictors of selective modulation and common signaling. **a)** Composition of minimal predictive models defined by machine learning. The predictive capacity of the model (y-axis) for a GR-mediated phenotype i.e. dependent variable (x-axis), is also indicated. See **Supplementary Table 2** for the full list of predictors. **b)** Linear regression (scatter plots) demonstrates their predictive power (*r*^2^), and its associated p-value. Each point represents the effects of a distinct ligand. **c)** Linear regression comparing the predictive power of the indicated peptide interactions for the indicated phenotypes and target genes. See **Supplementary Table 3** for *r*^2^ values. **d)** The Glut4-predictive power, *r*^*2*^, of target genes and peptide interactions observed with all compounds (**1-3** and **6-27**) were rank ordered, and then compared to the *r*^*2*^ observed within the C11-substituted (**13–20**) or C3-substituted (**6-12**) compound series. **e)** Linear regression demonstrating the Glut4-predictive power of *Bcl2l1* within the C11-substituted compounds series (**13–20**). Each datapoint represents the effects of a distinct C11-substituted compound on Glut4 translocation and *Bcl2l1* expression. See also **Methods**, **Supplementary Dataset**, and **Extended Data Fig. 3**.

This analysis revealed the degree to which target genes and interacting peptides functioned as selective predictors, or coordinated effects of GCs across the different biological processes (**Fig. 2a**–**b, Supplementary Table 2**, **Extended Data Fig. 3a–b**). We used linear regression to define the statistical power of individual predictors, where the r^2^ statistic (mathematically equivalent to the square of the Pearson correlation) defines the percent of variance in the dependent variable that is predicted by the independent variable. Ligand interaction patterns with GR and NCOR1 or NCOR2 were selective for predicting Synthesis, while NCOA1 and NCOA2 interaction profiles and expression of the *Bcl2l2* selectively predicted Glut4. In contrast, expression of *Sgk1* and *Socs2* were common regulators of both processes (**Fig. 2b**, **Extended Data Fig. 3b**). *Bcl2l1* encodes Bcl-xL, a mitochondrial, anti-apoptotic BCL-2 family protein that inhibits glucose metabolism and mitochondrial-mediated secretion of insulin by pancreatic β-cells ^23^, suggesting that it also contributes to glucose disposal in skeletal muscle.

Ligand-mediated NCOR2 peptide interaction with GR significantly predicted Synthesis, but not Glut4 or the top Glut4-predictive target genes (**Fig. 2c**, **Supplementary Table 3**). There was a robust connection between the coregulator peptides that predicted Glut4, showing very low *r*^*2*^ with the protein balance assays, but very high predictive power for the specific genes that best predicted Glut4 activity (**Fig. 2c**). While the MARCoNI assay is best viewed as a probe for ligand-regulated effects on surface structure, all of the peptides in **Figure 2c** represent bona fide nuclear receptor interaction motifs. Here we exploited the variance in ligand activity to identify common and selective transcriptional networks underlying GR-mediated phenotypes in skeletal muscle.

The variance in ligand activity profiles also allowed LCA to identify signal in what would typically be considered noise, such as the *Bcl2l1* and *Socs2* genes where most of the compounds had very little activity individually, but collectively proved highly predictive (**Fig. 2b, Extended Data Fig. 3ab**). Much of the gene expression data displayed an inflection point (**Extended Data Fig. 3c**). When we removed the data below the inflection point, *Fkbp5, Blc2l1*, *Tsc22d3* still significantly predicted Glut4 translocation (**Extended Data Fig. 3d**), suggesting that insulin-mediated glucose disposal can be fine-tuned by very small changes in target gene expression. We call these super-resolution analyses, similar to what we did with super-resolution X-ray crystallography, where comparing many structures bound to a series of compounds identified, within the noise of an individual crystal structure, ligand-induced perturbations that drive biological outcomes^8^. Thus, subtle changes in gene expression are sufficient to identify transcriptional networks underlying GR-mediated phenotypes using this approach.

We also found that substitutions on the steroidal scaffold that were designed to differentially perturb GR structure modulated distinct transcriptional networks. A comparison of ligand-dependent effects on Glut4 revealed a dramatic increase in predictive power by the C11-substituted compounds compared to the whole compound set, while the C3-substituted compounds exhibited a widespread reduction in *r*^*2*^ (**Fig. 2d–e**, **Extended Data Fig. 3e**, compare to **Fig. 2b**). This was evident despite their similar variances in the skeletal muscle profiling assays (**Extended Data Fig. 3f**), supporting the idea that C3- and C11-substituted compounds modulate GR-mediated phenotypes via different transcriptional networks and structural mechanisms, which we explore below with molecular dynamics simulations. This approach, LCA, demonstrates that looking at different classes of compounds, here comparing C3- and C11-substituted GCs, reveals different structural and signaling mechanisms to achieve their biological effects.

### Validation of individual predictors with gene perturbation studies

FKPB5 is a good candidate to coordinate metabolic effects of GCs at an organismal level, as it facilitates pAKT dephosphorylation by the phosphatase, PHLPP1^24^. In addition, *Fkbp5*-knockout mice show greater skeletal muscle insulin sensitivity and reduced adiposity on a high fat diet^25^. We electroporated *GFP* and *Fkbp5* or an atrophy-inducing control (*Foxo1*) expression plasmid into contralateral tibialis anterior (TA) muscles. The *Fkbp5-* or *Foxo1*-transduced muscles were approximately 10% smaller than control (**Fig. 3a**). Electroporation with *Foxo1* or *Fkbp5* inhibited insulin-induced *de novo* protein synthesis and pAKT *in vivo* (**Fig. 3b-c**, **Supplementary Fig. 1a**), demonstrating that *Fkbp5* regulates protein balance in skeletal muscle.

**Figure 3.**
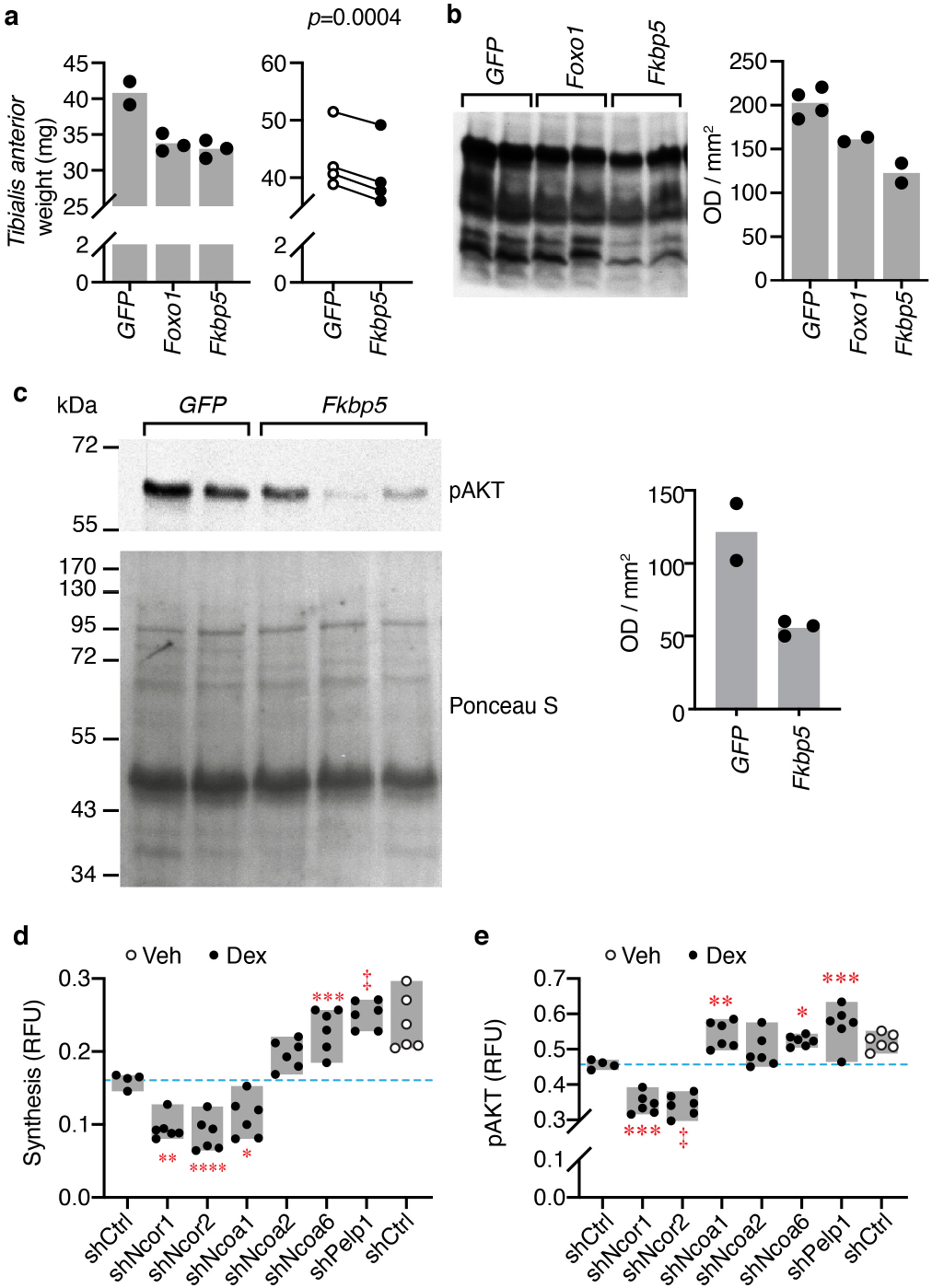
Functional validation of the predictive target gene, *Fkbp5*, and GR coregulators. **a**) Weight of Tibias Anterior (TA) muscles transduced with *GFP* (control), *Fkbp5,* or *Foxo1* genes in 2 independent experiments. Left panel, bars represent the mean, and n = 3 except for *GFP* where n = 2 TA muscles per condition. Right panel, each pair of datapoints represent TA muscles from the same mouse, n = 4 mice. Also see **Methods**. **b**–**c**) Whole lysates of transduced TA muscles were analyzed by Western blot and subsequent quantitation. **b**) In vivo SUnSET assay for Synthesis. Bars represent the mean; n = 2, except for *GFP* where n = 4 biologically independent samples. **c**) Insulin-induced pAKT levels. Bars represent the mean; for *GFP*, n = 2 and for *Fkbp5*, n = 3 biologically independent samples. Also see **Supplementary Fig. 1a**. **d**) SUnSET assay or **e)** in-cell Western for pAKT in C2C12 myotubes expressing the indicated shRNAs. Boxes represent the range; n = 6, except for Dex/shCtrl where n = 4 biologically independent samples. 1-way ANOVA, Sidak’s multiple comparisons test, adjusted p-values, \**p*_adj_ = 0.0235, \*\**p*_adj_ = 0.0018, \*\*\**p*_adj_ = 0.0007, \*\*\*\**p*_adj_ = 0.0004, ^‡^*p*_adj_ < 0.0001. **e**) Insulin-induced pAKT levels in C2C12 myotubes expressing the indicated shRNAs. Bars represent the range; n = 6, except for Dex/shCtrl where n = 4 biologically independent samples. 1-way ANOVA, Sidak’s multiple comparisons test, adjusted p-values, \**p*_adj_ = 0.0385, \*\**p*_adj_ = 0.0045, \*\*\**p*_adj_ = 0.0001, ^‡^*p*_adj_ < 0.0001. Also see **Supplementary Fig. 1b**.

Knockdown of *Ncor1* or *Ncor2* enhanced Dex-dependent inhibition of Synthesis (**Fig. 3d**, **Supplementary Fig. 1b**), consistent with the peptide binding data showing that GR interaction with NCOR1 and NCOR2 peptides were stimulated by anabolic compounds (**Fig. 2b**). In contrast, knockdown of *Ncoa6* and *Pelp1* fully reversed the inhibitory effect of Dex (**2**), while knockdown of *Ncoa22*, but not *Ncoa1*, partially reversed this effect (**Fig. 3d**, **Supplementary Fig. 1b-c**). In contrast, pAKT was regulated by *Ncoa1*, as well as *Pelp1* (**Fig. 3e**). These data support a model where GR recruits specific coregulators to distinct target gene sets that can have overlapping or different effects on skeletal muscle metabolism and protein balance.

### Selective GR modulators with improved activity profiles

We identified two promising GCs that selectively modulate inflammation and protein balance (**Fig. 4a–d**). Compounds **13-15** in the C11 series inhibited IL-6 with efficacies comparable to Dex (**2**) (**Extended Data Fig. 4a**). Among these, SR11466 (**15**) is a partial GR agonist (EC50 = 0.1 nM) that showed no effects on protein balance, and a slight improvement (i.e. less inhibition) compared to Dex (**2**) in the Glut4 assay (**Fig. 4a–c**, **Extended Data Fig. 4b-c**). We also identified SR16024 (**18**) as a full GR antagonist (IC50 = 1.4 nM) that stimulated Synthesis but did not suppress Degradation or increase Glut4 translocation (**Fig. 4b-c**, **Extended Data Fig. 4b-c**). **15** and **18** increased ψm, with the latter doubling ψm (**Fig. 4d, Extended Data Fig. 4d**).

**Figure 4.**
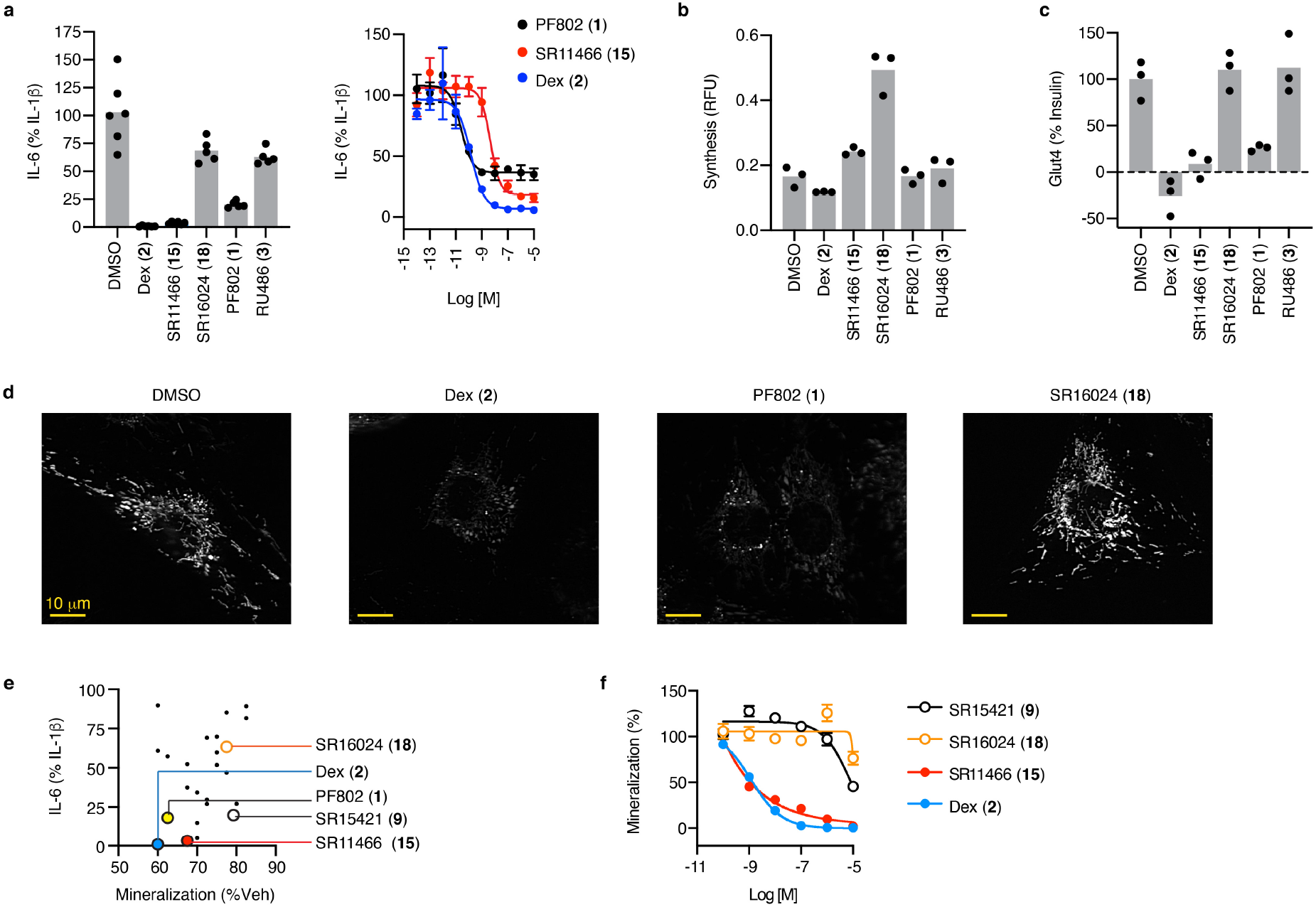
In vitro characterization of selective GR modulators with muscle-sparing activities. **a)** IL-1β-induced secretion of IL-6 by A549 cells treated with the indicated compounds were compared by AlphaLISA. Left panel bars represent the mean; right panel datapoints represent mean ± SEM; n = 3, except for the vehicle (DMSO) where n = 6 biologically independent samples. **b)** Effects of the indicated compounds on Synthesis in C2C12 myotubes were compared by SUnSET assay. Bars represent the mean; n = 3 biologically independent samples. **c)** SR16024 (**18**) does not inhibit myotube surface expression of Glut4. Effects of the indicated compounds on Glut4 in L6 myotubes. Bars represent the mean; n = 3 biologically independent samples. **d)** C2C12 myoblasts treated with the indicated compounds were stained with MitoTracker dye. Images are representative of 18 images per condition i.e. 2 independent experiments with similar results × 3 fields per well × 3 biologically independent wells per condition. **e)** The effects of all tested compounds on IL-1β-induced secretion of IL-6 by A549 cells (y-axis) and primary human osteoblast mineralization (x-axis). Datapoints represent the mean effects of a distinct compound, For IL-6, n = 3 and for mineralization, n = 4 biologically independent samples. **f)** Dose curves of indicated compounds in the mineralization assay. Data are mean ± SEM, n = 4 biologically independent samples. Also see **Extended Data Fig. 2**, **Extended Data Fig. 4,** and **Methods**.

As a further test for selective modulation, the compounds were profiled in primary human osteoblasts for effects on mineralization during 4 weeks of differentiation from mesenchymal stem cells. **15** showed a slight improvement in the inhibition of mineralization compared to Dex (**2**) and PF802 (**1**), while **18** showed a better profile (**Fig. 4e**). We also noticed SR15421 (**9**), a lower affinity compound with EC50 of 240 nM (**Extended Data Fig. 4c**), that was anti-inflammatory with less impact on mineralization (**Fig. 4e, Supplementary Fig. 2**). **9** is part of the C3-substituted series, where modifying the position of the methyl and the nitrogen on the methylpyridine had profound effects on inhibition of IL-6 secretion (**Fig. 4e**, **Supplementary Fig. 2**). Dose response curves reveal a moderately, but significantly improved effect of **15** on mineralization compared to Dex (**2**) (2-way ANOVA, drug p = 0.0018, drug × time p = 1.4 × 10^−^ 4), while **9** and **18** inhibited mineralization only at the 10 μM dose (**Fig. 4f**). The mineralization data showed very low correlations with the other assays, except for Synthesis (Pearson r = 0.54, p = 4 × 10^−3^), supporting the joint regulation of bone and muscle mass by GCs. These data demonstrate that it is possible to find GCs with improved bone- and muscle-sparing properties compared to Dex (**2**) using our in vitro profiling platform.

We opted to test the two high affinity ligands further in animal studies based on their favorable pharmacokinetics and on-target mechanism of action (**Fig. 5a–b**, **Extended Data Fig. 5a–f**, **Extended Data Fig. 6a, Supplementary Fig. 3**). In vivo, **15** strongly suppressed LPS-induced TNFα levels in the blood, while **18** was not inhibitory (**Fig. 5c**). We also assessed loss of lean mass following a larger dose of LPS, which was significantly worsened by Dex (**2**). Both **15** and **18** were significantly better than Dex (**2**), while **18** was protective against loss of lean mass compared to vehicle (**Fig. 5d**). The differential in vivo effects of **15** on inflammation and proteostasis demonstrate that it is a bona fide selective modulator, while **18** displayed stronger muscle sparing activities.

**Figure 5.**
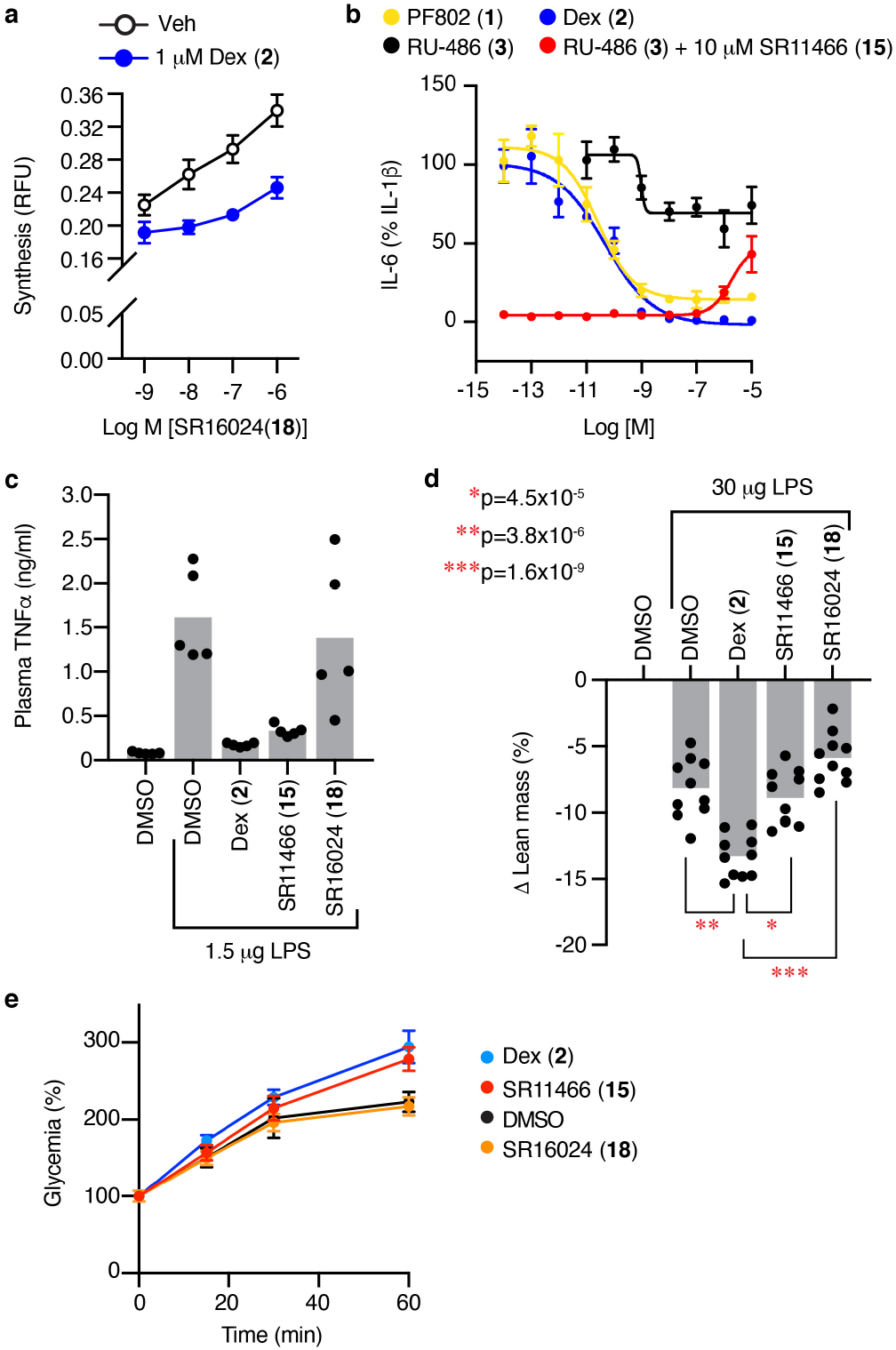
In vivo compound profiling and validation of on-target mechanism of action. **a)** Effect of the indicated compounds on Synthesis were compared by SUnSET assay in C2C12 myotubes. Datapoints are mean ± SEM, n = 4 biologically independent samples. **b)** IL-1β-induced IL-6 production by A549 cells treated with the indicated compounds was compared by AlphaLISA. Datapoints are mean ± SEM, n = 3 biologically independent samples. **c)** SR11466 (**15**) blocks the LPS-induced inflammatory response in mice. Plasma TNFα levels of mice treated with the indicated GR ligands (10 mg/kg Dex or SR16024, or 50 mg/kg SR1166) overnight before a 1-hr LPS challenge of 1.5 mg per mouse. Bars represent the mean, n = 5 mice per group. **d)** Changes in lean mass of mice treated as described in panel **c** were determined by whole-body NMR after an additional 18-hr LPS treatment of 30 mg per mouse. Bars represent the mean, n = 10 mice per group (5 × 2 experiments). 1-way ANOVA, Sidak’s multiple comparisons test, adjusted p-values, *p_adj_ = 4.5 ×10^−5^, **p_adj_ = 3.8 ×10^−6^, ***p_adj_ = 1.6 ×10^−9^. **e)** Glucose production was compared by lactate-tolerance test (LTT) after overnight fast. Data are mean ± SEM, n = 5 mice per group. See also **Extended Data Fig. 5 and Methods**.

A lactate tolerance test was then administered to probe for differences in liver gluconeogenesis following an overnight fast. Here again, Dex (**2**) caused significantly greater loss of lean mass after the fast; both **15** and **18** were significantly better than Dex (**2**) (**Extended Data Fig. 6b**); and **18** attenuated the loss of overall body weight from the fast (**Extended Data Fig. 6c**). The lactate tolerance test showed that Dex (**2**) and **15** increased the glucose production rate, while **18** did not (**Fig. 5e**), suggesting that selective GC effects on glucose uptake in skeletal muscle and gluconeogenesis in the liver may utilize common signaling mechanisms, such as ligand-selective coregulator recruitment, to integrate organismal stress responses.

### Structural features of GR linked to suppression of IL-6

C3 isomers with differences in the location of the methyl and nitrogen in the pyridine ring showed large differences in anti-inflammatory effects, demonstrating that the methylpyridinyl acetate binding under the AF-2 surface is an important allosteric regulator of GR. To understand the mechanism, we performed all atom molecular dynamics (MD) simulations comparingthe isomers **7** and **9**, which differ only in the placement of the methyl on the pyridine (**Fig. 1a**) but showed very different effects on suppression of IL-6 (**Supplementary Fig. 2**), where **9** suppresses inflammation but **7** does not. Molecular docking demonstrated that the compounds bound similarly to Dex (**2**), with the pyridine extending off the A-ring into a solvent channel, in between helix 3 (h3) and h5 (**Methods, Extended Data Fig. 7a**). In three independent 1 μs simulations of ligand-bound and ligand-free (apo) GR LBD, we observed no significant structural changes in the conformation of helix 12 (h12). However, ligand anti-inflammatory activity (Dex (**2**) > **9**>**7**> apo) was associated with a multi-Å decrease in the distance between the C-terminus of h11 and the N-terminus of h3, and a decrease in the heterogeneity of this h3/h11 interface (**Fig. 6a–c**). We also assessed patterns of correlated motion and found that the **7** showed less correlation between residues h3 and h11/h12 residues compared to Dex (**2**), with **9** again showing intermediate effects (**Extended Data Fig. 7b**).

We used dynamical network analysis^26^ to further probe for effects of the C3 substitutions on correlated motion between h12 and the solvent channel, by connecting pairs of nodes with their edges if they have satisfied a distance requirement (<4.5 Å) for ≥75% of the simulation time. Edge distance is inversely proportional to the pairwise correlations between two nodes; thus, a short path length indicates a strong correlation/communication. A pair of distal nodes are connected by the optimal (shortest) path and the suboptimal (longer) paths with the length defined as the sum of edge distances along the path. We analyzed the top 1,000 suboptimal paths between residue Glu755 and Arg614 as two nodes to study the allosteric communication between h12 and the C-terminus of h5, where the modifications on the steroidal A ring extend to the solvent channel. With apo GR, this communication was through h11 and along h5, while with Dex (**2**) it wrapped around the ligand-binding pocket from h12 to h3 and then through the β-sheet (**Fig. 6d**), enabled by the Dex-induced closing of the distance between h3 and h11 (**Fig. 6b**). With Dex (**2**), this was associated with much longer path lengths (**Fig. 6e**), representing overall weaker allosteric communication. With **9**, there were less of these extended path lengths, while **7** more closely resembled the apo GR (**Fig. 6d-e**). An analysis of residues contacting the C3-substituted pyridine showed that differential positioning of the Arg611 side chain on h5, which can form h-bonds with the pyridine or A-ring ketone of Dex (**2**), can drive the orientation of the steroid core to control the dynamics and surface structure and associated receptor activity (**Fig. 6f, Extended Data Fig. 7c–e**).

**Figure 6.**
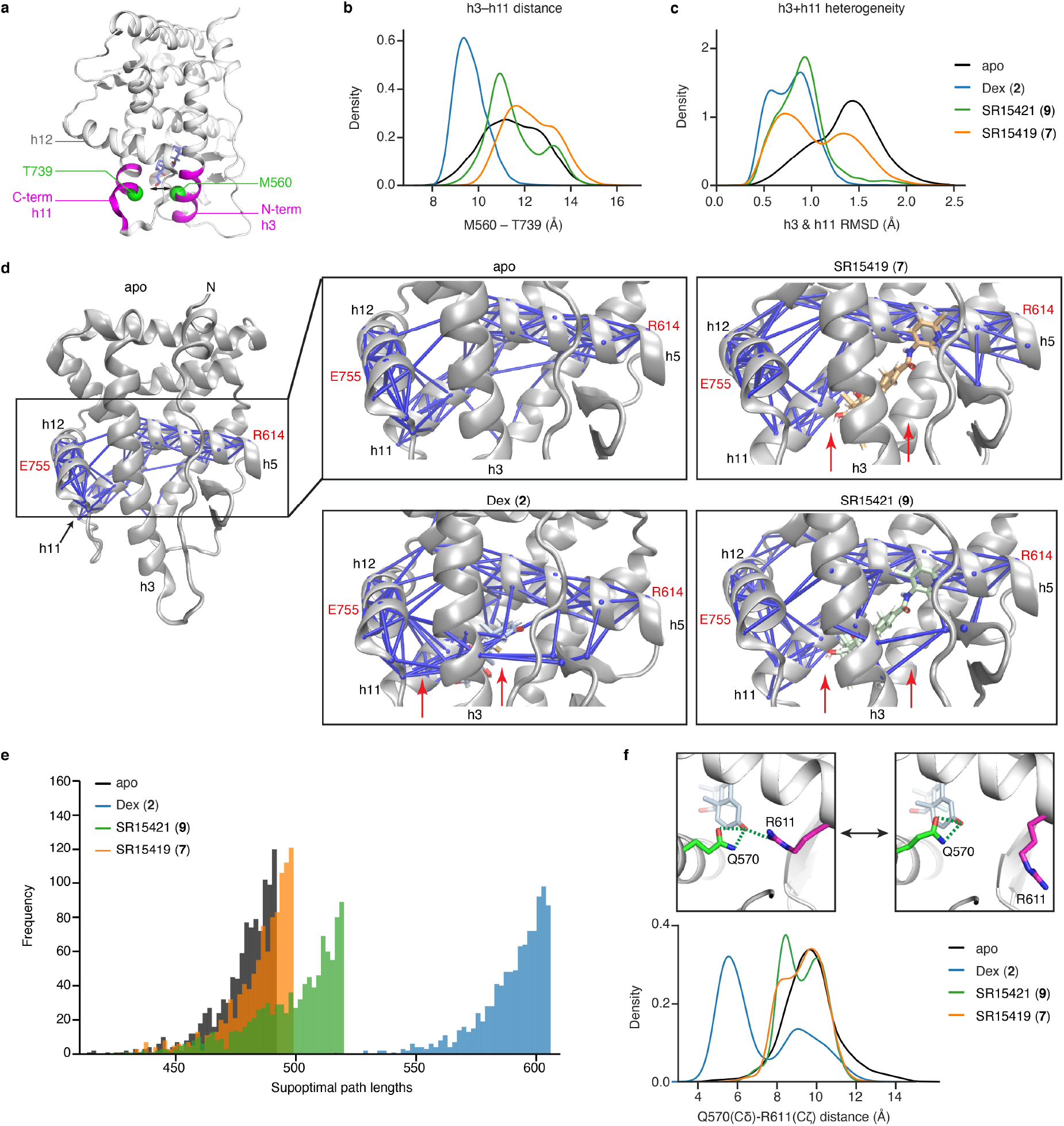
C3 substitutions in the steroid scaffold alter allosteric communication between ligand and activity. **a)** Ribbon diagram of the GR-LBD illustrates the C-terminus of helix 11 (h11) and N-terminus of h3 where we observed ligand-induced conformational effects in three 1 μs molecular dynamics simulations. **b)** Distance distribution plots of the Cα distance between Met560 (h3) and Thr739 (h11). **c)** Backbone Cα, C’, N, and O RMSD distribution plots of the h3 and h11 regions colored magenta in (**a**). **d)** Dynamical network analysis of suboptimal pathways for correlated motion between E755 and R614. Red arrows indicate pathways found with Dex-bound GR. **e)** Histogram showing the suboptimal pathlengths with the indicated ligands. **f)** Distance distribution plots of Q570 (Cδ) and R611 (Cζ) side chain atom distances as a proxy to determine the relative populations inward R611 conformations that can interact with ligand. Inserts show R611 inward (left) and outward (right) side chain conformations extracted from the Dex-bound simulations.

## Discussion

We used structure-based design and LCA to reveal how GCs modulate different transcriptional networks (coregulators and target genes) to physiologically control glucose disposal and protein balance during acute nutrient deprivation, and how different classes of these ligands with substitutions at C3 or C11 utilize these signaling pathways. By examining closely related compounds, we determined that protein balance and Ψm were intercorrelated but not correlated with effects on Glut4 translocation and suppression of IL-6. Using a physiologically relevant skeletal muscle profiling platform optimized to study insulin signaling during nutrient deprivation, we identified SR11466 (**15**) as a partial agonist that was not catabolic, demonstrating in vivo selective modulation of anti-inflammatory activity. **15** was slightly anabolic for protein balance, while maintaining strong anti-inflammatory activity in vivo. Such compounds may be useful treatments for cachexia, muscular dystrophies, back pain or osteoarthritis. We also identified SR16024 (**18**) as a highly anabolic GC that inhibits fasting- and LPS-induced weight loss, while stimulating Ψm and protein synthesis in response to insulin. Further studies are needed to evaluate the long-term effects of **18**, but GR antagonists have been tested clinically for depression^27^, and may have efficacy in muscular dystrophies and enzalutamide-resistant prostate cancers that have switched from androgen to glucocorticoid dependency^28^. While preliminary, the lack of SR15421 (**9**) effects on mineralization suggests that it may also be possible reduce the osteoporotic effects of agonist GCs using physiologically relevant profiling assays.

The unbiased nature of LCA enabled the identification of a number of unexpected signaling and biophysical properties of GR. Insulin receptor signaling pathways control anabolic effects and glucose disposal, which can be disconnected by transcriptional control of pathway-selective regulators, often with subtle changes in gene expression. Covariance among assays reflects common underlying signaling mechanisms driven by common receptor conformations, coregulators, and target genes. The correlation between effects on bone and skeletal muscle was not surprising, but the connection between anti-inflammatory and metabolic effects of GCs in different cell types was unexpected and may point to an organismal level of evolutionary connection between underlying signaling pathways. GC coordination of the inflammatory response often coincides with immune differentiation^29,30^, which requires metabolic adaptations 31,32. This suggests that GCs may use a common set of coregulators to coordinate anti-inflammatory effects with modulation of metabolism. LCA provides a mechanism to identify common signaling mechanisms among different cell types and processes and how they are utilized by structurally distinct classes of ligands.

GC-regulated transcription is controlled via underlying biophysical properties of GR, such as oligomerization, nuclear translocation, and on/off rates of interactions with an ensemble of response elements, coregulators, and collaborating transcription factors. We have shown here that LCA predicts that ligand-selective surface conformers—identified from a structural peptide interaction assay—can predict differential effects of the ligands, which we validated with knockdown of selected coregulators. We and others have observed a similar dichotomy where NCOA2 is required for NF-κB-induced cytokine production, but then switches to a repressor upon recruitment of GR to the promoter^33,34^. This is representative of a more general phenomena of receptors using many coregulators to regulate a single gene, but also some shared and some differential coregulator usage across genes^14,35,36^. The selective association of GR nuclear translocation with glucose disposal and not protein balance was also surprising. This suggests that GCs modulate insulin dependent Glut4 translocation via a graded transcriptional program that is highly sensitive to the amount of nuclear GR, and control protein balance through transcriptional responses that require a threshold amount of nuclear GR. It is remarkable that changing the position of a single methyl in the solvent channel drove such divergent effects of the C3 compounds on inflammation, while the molecular dynamics simulation revealed this region to be an important allosteric regulator of GR structure and function. Collectively our finding show that complex biology can be simulated by a set of simple biophysical models^9,37^ and their importance identified with LCA.

## Acknowledgements

N.E.B. was supported by the BallenIsles Men’s Golf Association. K.W.N. was supported by G.S. Humane Corp. W.M. Keck Foundation Grant to EAO.

## Author contributions

Conceptualization, N.E.B. and K.W.N.; Methodology, N.E.B., K.W.N., S.V.B, J.C.N., G.L.H., M.C., R.H.; Investigation, N.E.B., S.S., S.V.B., J.C.N., C.C.N., D.A.S., R.H., D.J.K., X.L., O.E., Z.J., T.M.K.; Resources, Z.J., T.M.K.; Writing – Original Draft, K.W.N., N.E.B., J.C.N.; Writing–revision, K.W.N., J.C.N., X.L., E.A.O., D.J.K, E.O.; Supervision, K.W.N., N.E.B., G.L.H., T.M.K., E.A.O., T.I.

**Extended Data Fig. 1.**
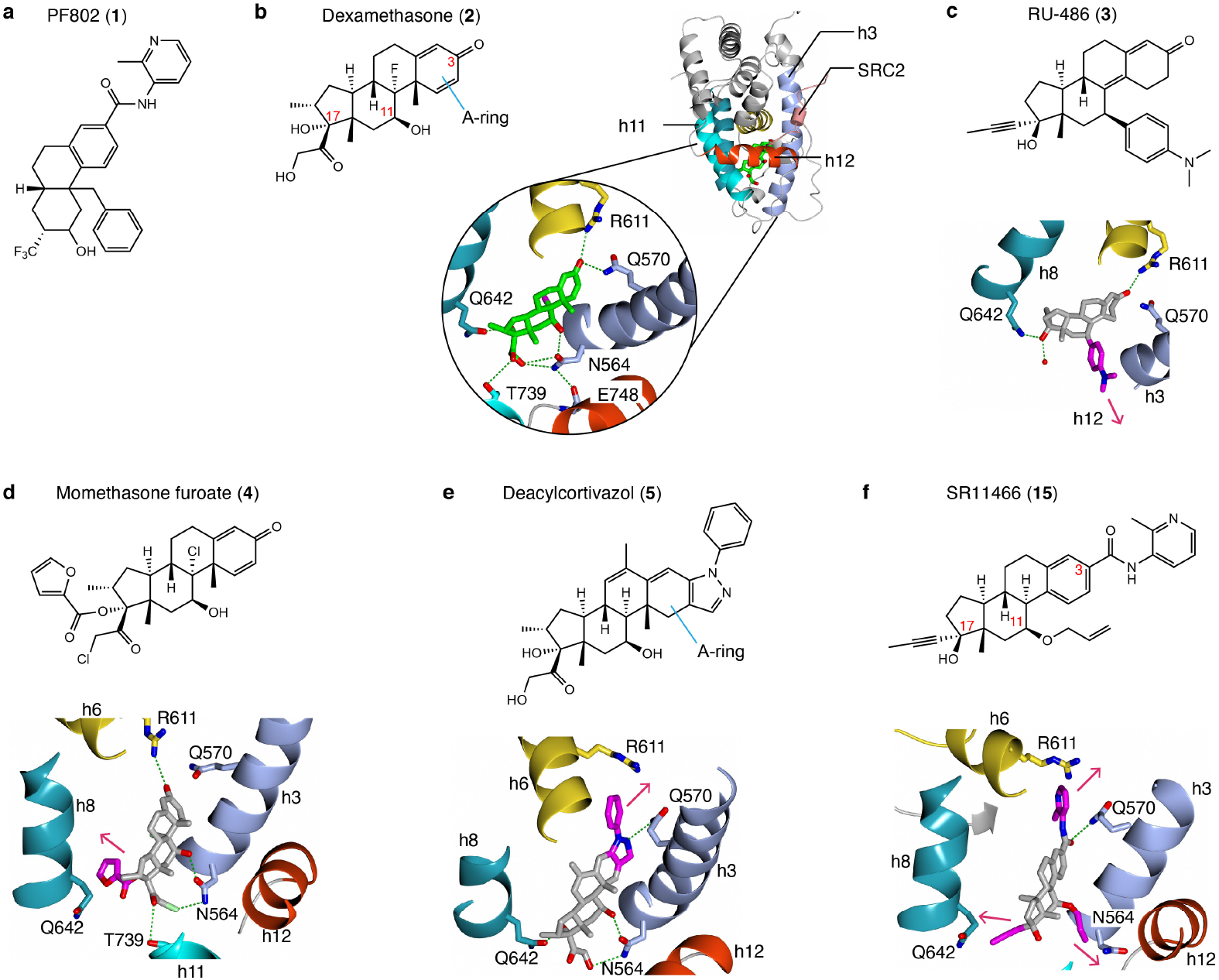
Structure-based design of GR ligands. **a)** Chemical structure of the selective GR modulator, PF802. **b)** Crystal structure of the dexamethasone (Dex) bound GR ligand-binding domain (LBD). Helix 12 is colored red and the NCOA2 coregulator peptide binding in the AF2 binding surface is colored coral. Carbons-3, −11, and −17 are indicated in the chemical structure (1M2Z.pdb). **c)** The bulky dimethylanaline group attached at C11 in RU-486 displaces h12 from the agonist position to disrupt the AF2 surface and generate antagonism (1NHZ.pdb). **d)** Substitutions at C3, as seen with the furoate group in momethasone furoate or the propyne in RU-486 target a small internal pocket to increase affinity (4P6W.pdb). **e)** Substitutions at C3 of the steroidal A-ring enter the solvent channel underneath the AF2 surface, potentially changing the shape of the surface and the ensemble of interacting coregulators (3BQD.pdb). **f)** Model of SR11466 (**15**) bound to the GR LBD

**Extended Data Fig. 2.**
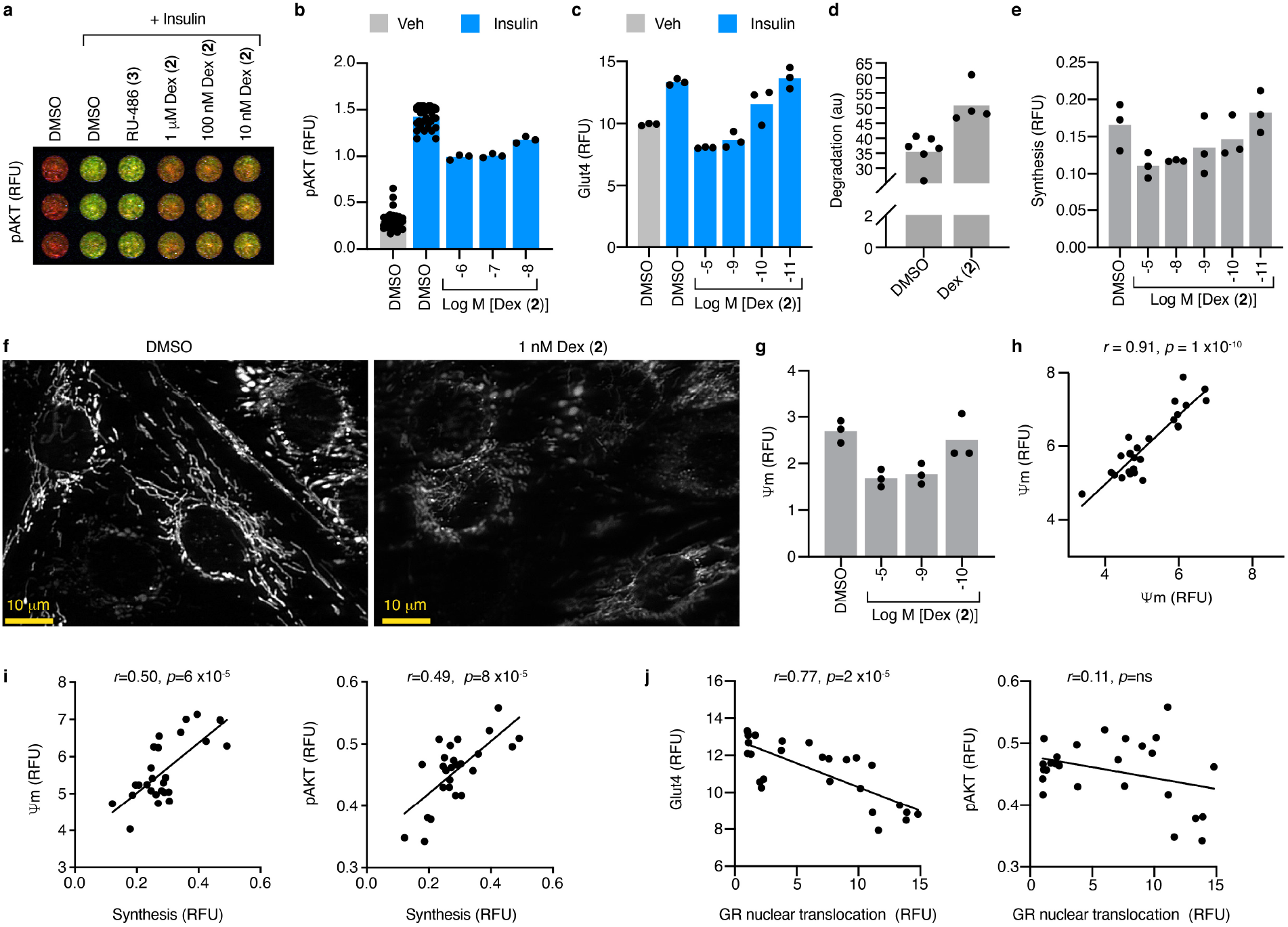
Quantitative phenotyping assays for GC action in skeletal muscle. **a-e**) Myotubes were nutrient-deprived, pre-treated with DMSO, RU-486, or Dex, and treated with insulin as outlined in **Methods**. **a)** Effect of insulin on pAKT levels in C2C12 myotubes were compared by In-Cell Western assay (ICW) 48 h after treatment with RU-486 or Dex. **b)** Quantitation of pAKT in C2C12 myotubes compared by ICW. Bars represent the mean; n = 3, except for DMSO where n = 36 biologically independent samples. **c)** ICW for surface expression of Glut4 on L6 myotubes. Bars represent the mean; n = 3 biologically independent samples. **d)** C2C12 myotubes were assayed for protein degradation by release of tritiated phenylalanine. Bars represent the mean; for DMSO, n = 6, and for Dex, n = 4 biologically independent samples. **e)** ICW for protein synthesis by insulin-induced incorporation of puromycin into C2C12 myotube surface proteins. Bars represent the mean; n = 3 biologically independent samples. **f-g**) High-content imaging and analysis of C2C12 myoblasts stained with MitoTracker™ dye. Images are representative of 18 images per condition; i.e. 3 fields × 3 biologically independent samples per condition in each of 2 independent experiments. **h**) Assay reproducibility from screening 22 compounds on two separate occasions. The Pearson correlation coefficient, r, and its associated p-value is indicated. Each datapoint represents the mean effect of a distinct compound. Also see **Methods**. **i**–**j**) Linear regression demonstrates the predictive power (*r*^2^), and associated p-value for the indicated variables. **i**) Ψm and pAKT predict Synthesis (p = 6 ×10^−5^ and 8 ×10^−5^, respectively). **j**) GR nuclear translocation selectively predicts Glut4 (p = 2 ×10^−5^) but not pAKT. Each datapoint represents the effects of a distinct ligand.

**Extended Data Fig. 3.**
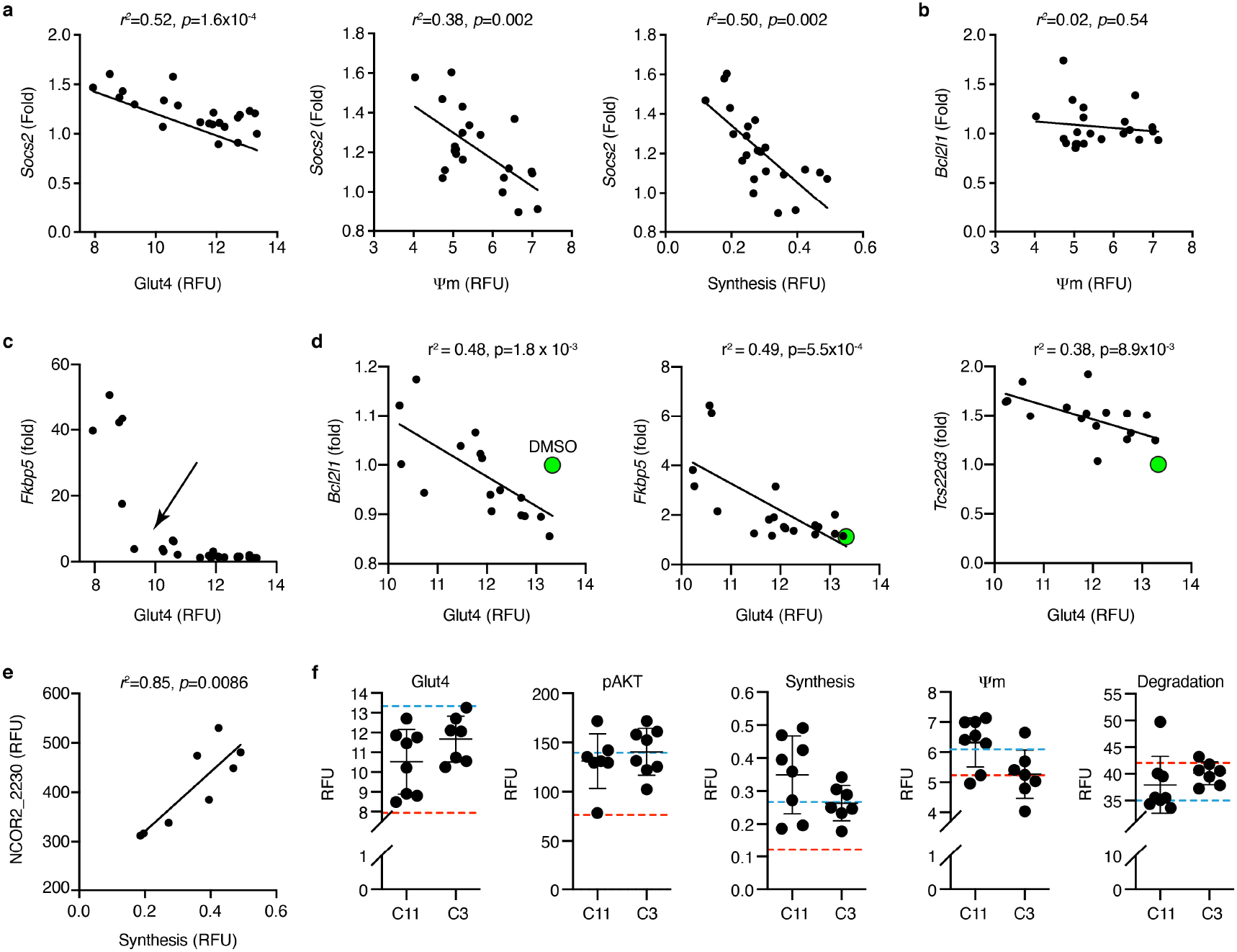
Relationships among specific genes, peptide interaction assays, and GR-mediated phenotypes. **a-e**) Linear regression was performed for the indicated assay pairs, where each point represents a different compound. **a)** Effects of the ligands on *Socs2* expression predicts Glut4 translocation, ψm, and insulin-stimulated protein synthesis. **b)** Ligand-dependent expression of *Bcl2l1*, which encodes the mitochondrial anti-apoptotic protein, Bcl-xL does not predict effects of on ψm. **c)** *Fkbp5* expression as a predictor of Glut4 translocation shows an inflection point (arrow). **d)** The Glut4 data was truncated below the inflection point shown in **c**). **e)** GR interaction with an NCOR2 peptide predicts protein synthesis in the C11 subset of ligands. **f)** The C11- and C3-substituted compounds showed similar variance in the skeletal muscle profiling assays. Blue dashed line, vehicle; red dashed line, Dex. Each datapoint represents the mean effect of a distinct compound, n=3 biologically independent samples. The error bars represent the mean ± SD of each compound series. For C11, n=8 distinct compounds; for C3, n=7 distinct compounds. See also **Fig. 2a-e**

**Extended Data Fig. 4.**
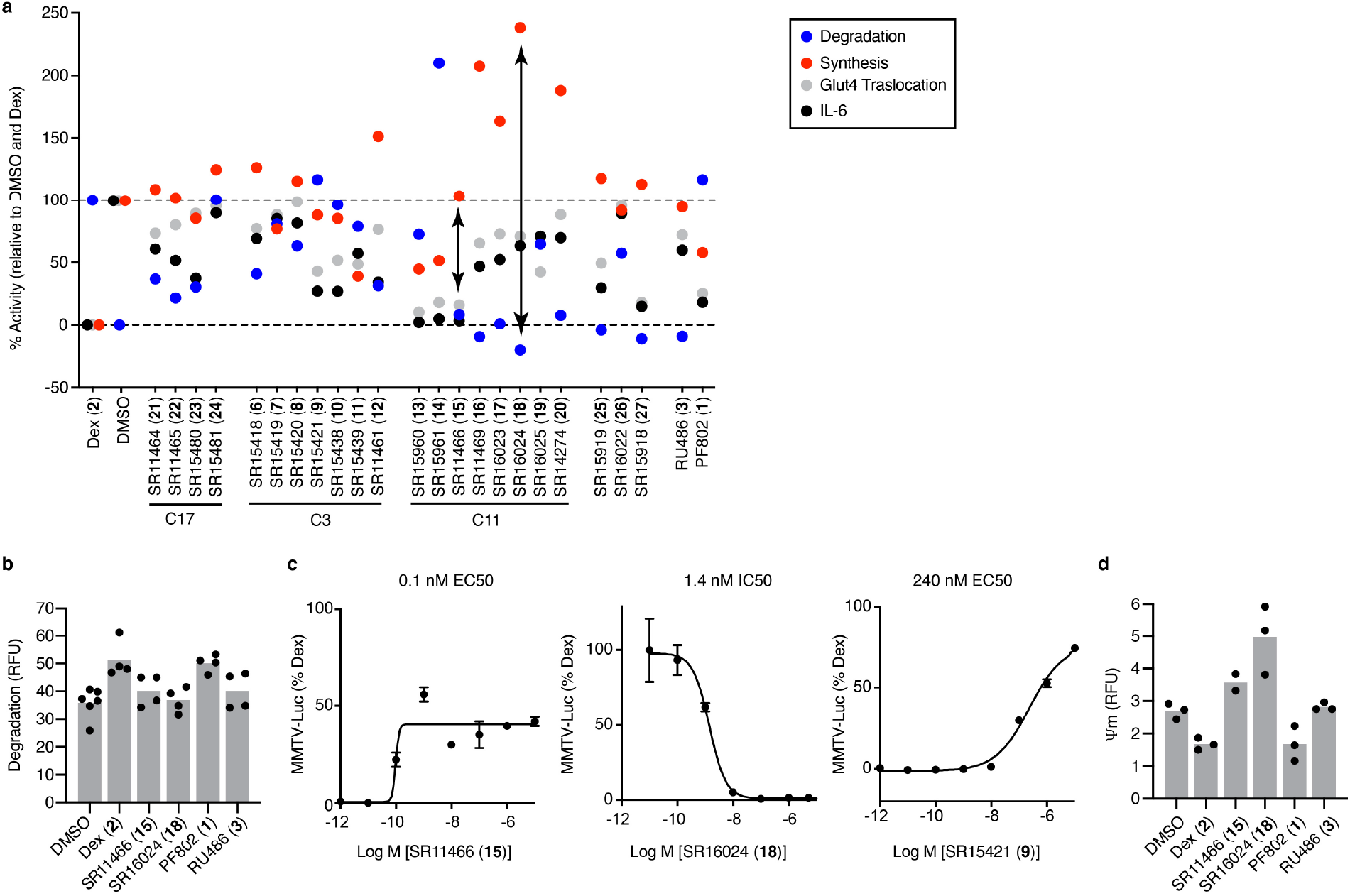
Compound structure-activity relationships. **a)** Individual compound data for protein degradation, insulin-stimulated protein synthesis, and Glut4 translocation in myotubes, as well as effects on IL-1β-stimulated secretion of IL-6 by A549 cells. Lead compounds are indicated with arrows. Among the 3 compounds with full suppression of IL-6 (**13,14,15**), only **15** did not inhibit protein synthesis or stimulate protein degradation. **18** showed the greatest anabolic effects, with stimulation of protein synthesis and inhibition of protein degradation. **b)** Protein degradation in myotubes assayed as described in **Extended Data Fig. 2** and **Methods**. Bars represent the mean, n = 4, except for DMSO where n = 6 biologically independent samples. **c)** 293T cells were co-transfected with a GR expression plasmid and MMTV-luciferase reporter. The next day cells were treated with the indicated compounds for 24 h and probed for luciferase activity. For SR16024, was the cells were cotreated with 1 nM Dex. Data are mean ± SEM, n = 3 biologically independent samples. **d)** Mitochondrial potential of myotubes assayed as described in **Extended Data Fig. 2** and **Methods**. Bars represent the mean; n = 3, except for **15** where n = 2 biologically independent samples.

**Extended Data Fig. 5.**
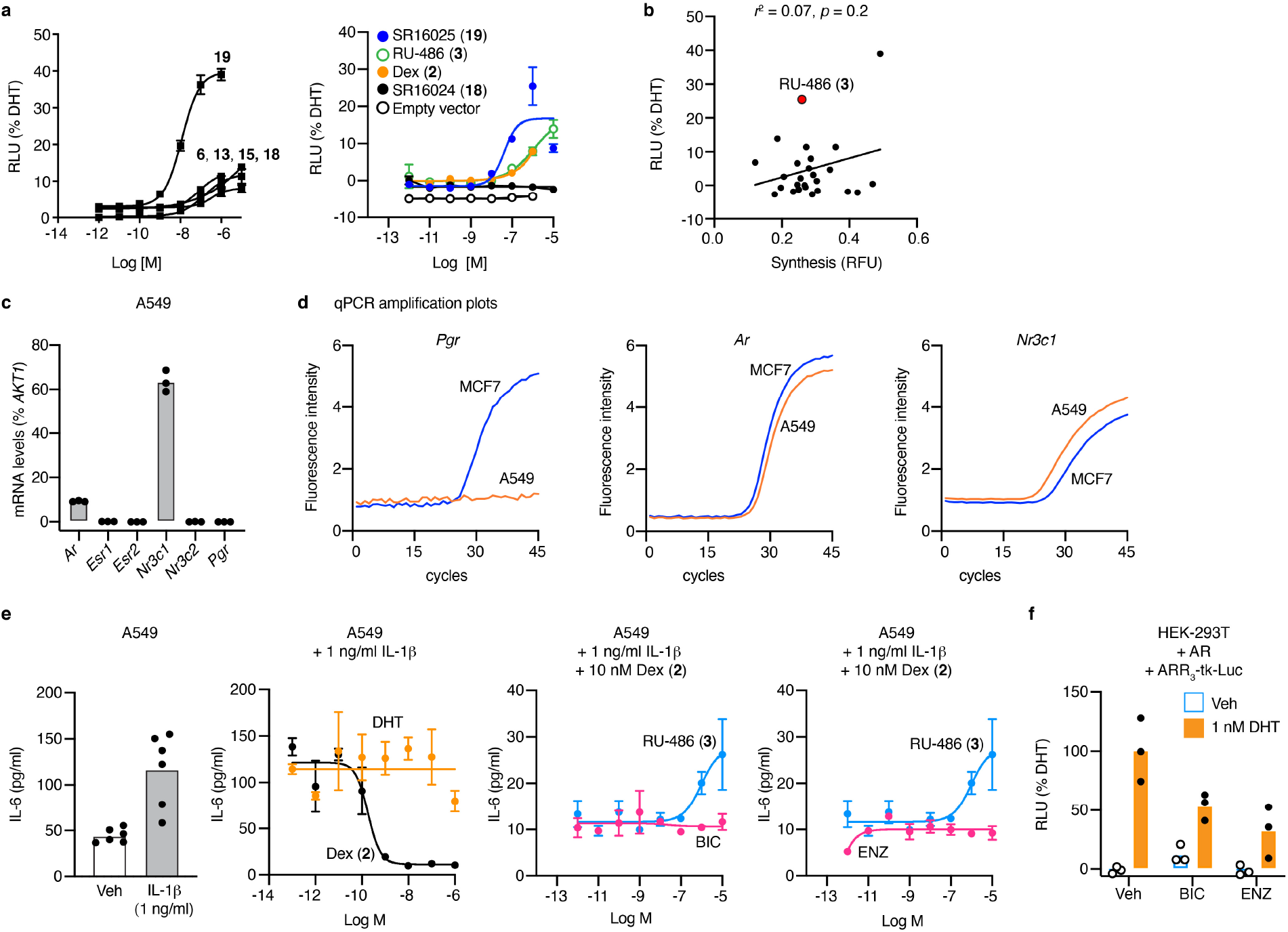
On-target mechanism of action studies. **a)** Reporter activity in steroid-deprived 293T cells co-transfected with an androgen-responsive ARR_3_-tk-luc reporter and an androgen receptor (AR) expression plasmid or empty vector control, and then treated with the indicated compounds for 24 h. Dose curves for compounds that stimulated AR activity (left) and the indicated compounds (right) are shown. None of the compounds showed activity with the empty vector control. **18** and **19** are isomers differing only in the position of the chlorine on the benzyl substitution. Datapoints are mean ± SEM; n = 3 biologically independent samples. **b)** Linear regression demonstrating that ligand-specific AR activity profiles do not correlate with protein synthesis. **c)** Expression of steroid receptor mRNAs in A549 cells. Only *Ar* which encodes AR, and *Nr3c1* which encodes GR were detected by qPCR. Bars represent the mean; n = 3 independent samples. Also see **Supplementary Fig. 3**. **d)** Representative qPCR amplification plots for *Pgr*, *Ar*, and *Nr3c1* in A549 versus MCF7 cells. *Pgr*, which encodes the progesterone receptor, is not expressed in A549 cells. **e)** AR antagonists do not reverse the effects of Dex on IL-6 secretion. IL-6 levels in A549 cell media were measured by AlphaLISA after overnight exposure to the indicated conditions. DHT, 5α-dihydrotestosterone; BIC, bicalutamide; ENZ, enzalutamide. For the controls (left), bars represent the mean; n = 6 biologically independent samples. For dose curves, datapoints are mean ± SEM; n = 3 biologically independent samples. **f)** Luciferase assay showing the effects of 1 nM DHT, 1 μM BIC and 1 μM ENZ on AR activity, demonstrating that the antagonists have cellular activity. Bars represent the mean; n = 3 biologically independent samples.

**Extended Data Fig. 6.**
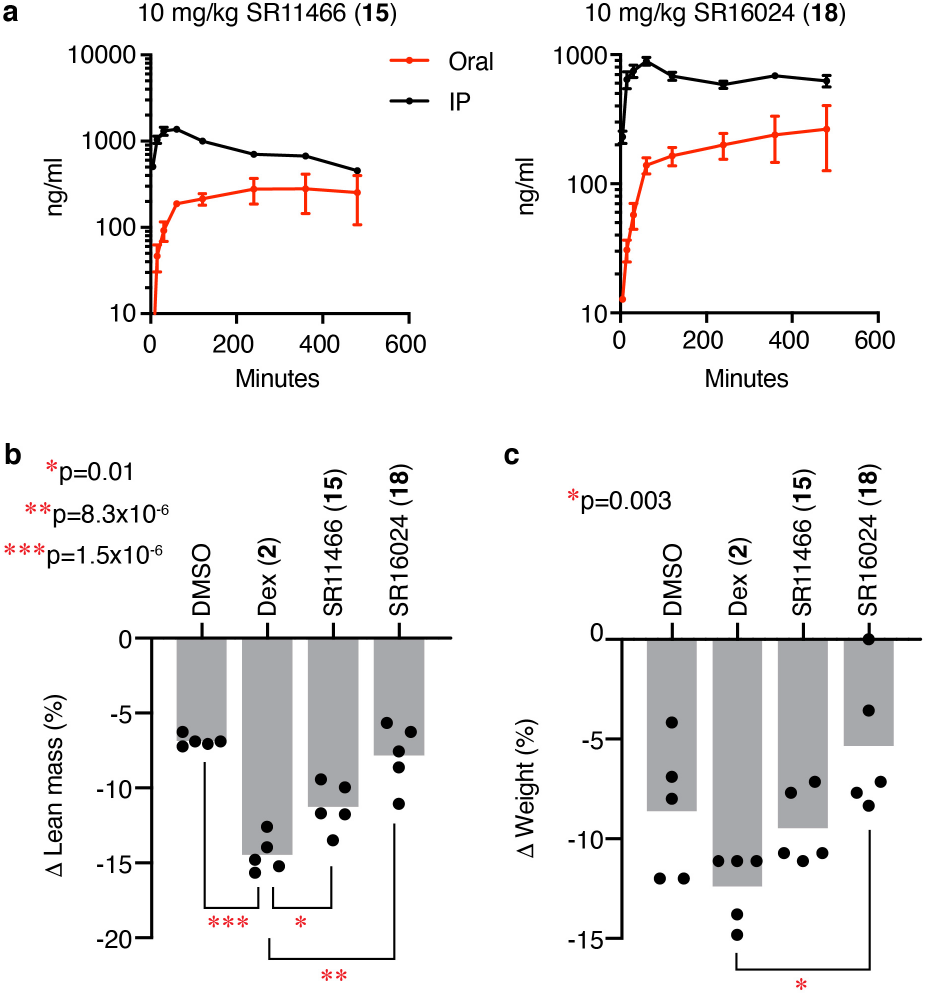
In vivo compound profiling. **a**) Mouse pharmacokinetics studies of the indicated compounds. Data are mean ± SEM; n = 3 biologically independent samples. **b-c**) Changes in the lean mass and body weights of male C57BL/6 mice treated with (10 mg/kg Dex or SR16024, or 50 mg/kg SR11466) and fasted overnight. Bars represent the mean; n = 5 mice per group (in each of 2 independent experiments). 1-way ANOVA, Sidak’s multiple comparisons test, adjusted p-values are indicated.

**Extended Data Fig. 7.**
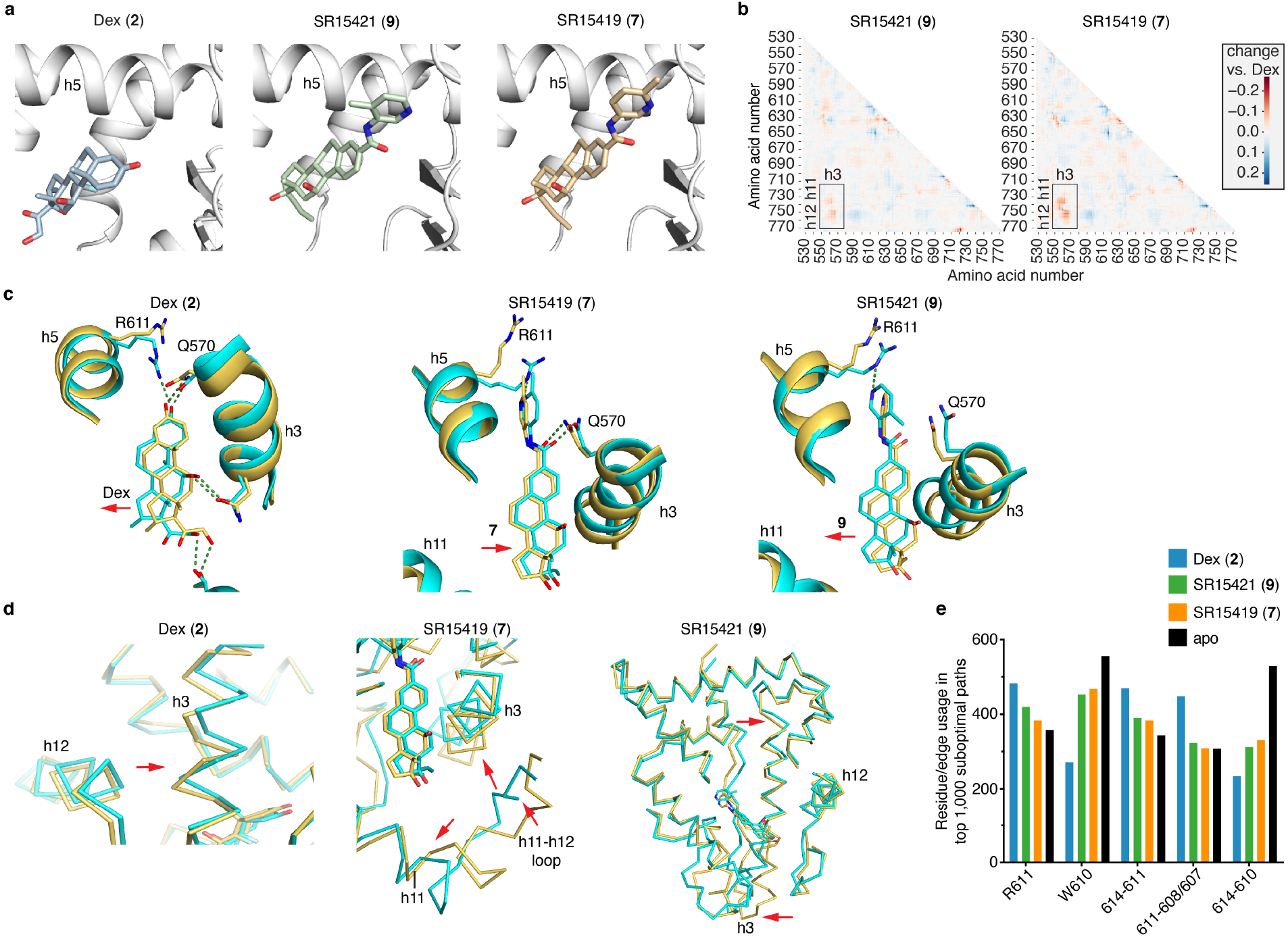
Docking and molecular dynamics simulations. **a)** Ribbon diagram of GR LBD bound to the indicated ligands. **7** and **9** were docked with Autodock Vina. **b)** Differential analysis of correlated motion between Cα atoms from the simulations with the indicated ligands subtracted from Dex. **c)** Formation of a hydrogen bond with R611 differentially shifts the position of the ligands. **d)** Formation of the hydrogen bond R611-induced changes in surface structure (red arrows). With Dex, there was a shift in h12 and the C-terminus of h3. With **7**, the C-terminus of h11 and N-terminus of h3 were shifted further apart, and away from h12. This destabilization of the h12 interface with h3 and h11 explains why this compound is an antagonist, a mechanism we have called “indirect antagonism.” With **9**, there was a rotation of both ends of h3. **e)** Usage of amino acid residue and edge in the suboptimal pathways between h12 E755 and h5 R614, demonstrating that Dex preferentially utilized R611 instead of W610 as a pathway for correlated motion. Also see **Methods.**

