## Supplementary Information for "Chemical Systems Biology Reveals Mechanisms of Glucocorticoid Receptor Signaling"

### Online Methods

#### Quantification and statistical analyses

All statistics were done using GraphPad Prism unless otherwise stated. Random Forest/Boruta were done with R. For Pearson correlation, the p-value (two-tail) tested the null hypothesis that the true population correlation coefficient for that pair of variables is zero. Two-tailed unpaired Student's t-test was done to compare only two groups. One-way or two-way ANOVA was used for comparisons of more than two groups with correction for multiple hypothesis testing. All values were expressed as the mean ± S.E.M. The compound injections were done by one person and the determination of weight/body composition done by another person blinded to the treatment groups. For the high content imaging assay of myotube diameter and the in cell western blots, images were visually inspected in a blinded fashion for the presence of intense foci or speckles and those images were not included in the analyses. For the IL-6 dose curves, data were identified through the linear regression function in GraphPad Prism.

***Machine learning:*** Each Boruta (version 6.0)[22] and *randomForest* package (version 4.6)[21] predictionwas performed with three different random seeds on Z score transformed data. Variables that were confirmed in 2 or more runs were selected as predictors. Sample *R* scripts are available at: https://github.com/jnwachuk/ML_in_GR_signaling_networks.

#### Nascent RNA-seq

$2x10^6$ C2C12 myoblasts were grown in 15 cm dishes to 90% confluency and differentiated in media with 2% steroid-free FBS. After 5 days, myotubes are treated with 1 uM Dex or vehicle, in duplicate plates, for 2 hr. Nuclei were isolated, washed, and lysed with a 10% NP40 buffer and centrifuged. The pellet was washed 2x with TRIzol™ reagent (Invitrogen) to remove unbound RNA. The chromatin pellet was then dissolved in Trizol at 65°C and RNA was isolated using standard protocols (e.g. Qiashredder and RNAEasy column purification). Library preparation was performed using TruSeq stranded total RNA kit (Illumina) and sequenced on NextSeq500 (Illumina). A total of 30-40 million 2x75bp paired-end mapped reads were obtained per sample. The sequencing reads (fastq files) were mapped to the mm9 genome using Tophat [38]. The number of reads falling into each gene defined in the RefSeq gene annotations was quantified using HTSeq-count [39]. The DESeq software [39] was used to detect DEGs.

**Nanostring nCounter gene expression analyses**

Myoblasts were differentiated for 5 days and then switched to charcoal stripped serum for 24 hr, and then treated with 1 μM GCs for 6 hrs. RNA was isolated with RNeasy (Qiagen). Gene expression analyses were performed using the manufacturer's instructions.

**Animal studies**

All animal procedures were approved by the Scripps Research Institute IACUC. Male C57BL/6J mice, 8–10 wk old, were acquired from the Jackson Laboratories (Bar Harbor, ME) and used for all conditions. Mice were housed under a 12-h light/dark cycle with *ad libitum* access to food and water unless otherwise stated. Before all surgical procedures, mice were anesthetized with gaseous isoflurane (Baxter, USP, USA). Before tissue extraction, the mice were euthanized with $CO_2$ and cervical dislocation.

**LPS-induced muscle atrophy:** Mice were treated with 1.5 μg LPS (Sigma L2880) for 1 hr and then blood drawn for determination of TNFα (ELISA, R&D Systems, MTA00B). Mice were injected with 30 μg LPS. Body composition was determined after 23 hrs using time domain nuclear magnetic resonance (TD-NMR) in a Bruker minispec.

**Lactate tolerance test:** Mice were administered GCs at 10 mg/kg IP and fasted overnight. The next morning, lactate (1.5 g/kg) was injected intraperitoneally and blood glucose assayed with Accu-check glucose monitor .

**Electric-pulse-mediated gene transfer and in vivo SUnSET assay:** Plasmid DNA (50 μg) was transfected into mouse tibialis anterior (TA) muscles by electroporation as previously described[42]. 1 hr before electroporation mice were anesthetized and a small incision was made through the skin and then injected with 30 ul of 0.5 U/Al hyaluronidase and plasmid DNA. Plasmids (Expression Ready MGC cDNA Library) were purified with an EndoFree plasmid kit (Qiagen, Valencia, CA, USA), and resuspended in 71 mM sterile PBS. 7 days later, mice were anesthetized and injected i.p. with 0.040 mol/g puromycin dissolved in 100 μl of PBS. 30 min after injection, muscles were extracted and frozen in liquid $N_2$.

**Western blot analysis**

Whole cell extracts were prepared in 20 mM HEPES, pH 7.4, 1% TX-100,1x phosphoSTOP complete inhibition cocktail tablets [Roche], and 1X Complete EDTA-Free protease inhibitor cocktail [Roche]). Frozen muscle tissue was homogenized in lysis buffer T-PER® Tissue Protein Extraction reagent (Thermo Scientific) by a polytron homogenizer and sonication. Protein concentration was determined using the protein assay reagent (Bio-Rad). Sixty to eighty μg of protein was separated by SDS-PAGE and transferred to nitrocellulose membranes for immunoblotting.

**Cell-based assays**

C2C12 myoblasts, L6 myoblasts, and A549 cells (ATCC) were cultured in high glucose DMEM supplemented with glutamine and 10% FBS. For steroid-free culture conditions, cells were grown in charcoal:dextran-stripped FBS (Gemini Bio-products, cat no. 100-119). All assays were performed in C2C12 cells (ATCC) except for the Glut4 translocation assay, which was performed in rat L6 myocytes.

**pAKT:** C2C12 myoblasts were seeded at 70% confluency and 48 hrs latter washed with PBS and switch to differentiation media consisting of phenol-free DMEM supplemented with glutamine and charcoal stripped serum (Sigma 18l289). Partial media changes were performed every 48 hrs to prevent lifting of the tubes from the wells. After six days in differentiation media, the cells were serum starved, treated with ligands. The next day, cells were stimulated with 100 nM insulin for 30 min., fixed in 3.7% formaldehyde for 20 min, permeabilized with TBS + 0.1% Triton X-100, washed in TBS and incubated for 2 h in blocking solution (Rockland MB-070), then overnight in primary antibodies at 4°C. Then the plates were washed 5 times for 5 minutes with TBS-T and then incubated with secondary antibodies for 1 hr wash and analyzed using a LI-COR Odyssey imaging system.

**Glut4 translocation:** Low passage, L6 rat myoblast (ATCC), were cultured in high glucose DMEM supplemented with glutamine and 10% serum (proliferation media). On day 1 of differentiation, 2,500 cells were seeded on 384-well plates in 50 μl of proliferation media that was replaced every other day. On day 5 cells were washed and replaced with differentiation media, which was then replaced daily. After 3 days of differentiation the cells were ligand treated and serum starved overnight. Next day we replace the medium with 100 µl DMEM ± 1 µM insulin (Sigma I0516) without serum or glucose and incubate for 1 hour and then fixed with 3.7% formaldehyde for 20 minutes. Non-permeabilized cell were then washed in TBS and blocked for 2 hours (Rockland MB-070) then incubated in primary anti-Glut-4 antibody in blocking buffer for 48 hours at 4 °C. Then the plates were washed 5 times for 5 minutes with TBS-T and then incubated with secondary antibody for 1 hr wash and analyzed using a LI-COR Odyssey imaging system.

**Protein synthesis (in vitro SUnSET assay):** C2C12 myoblasts were cultured and differentiated in 384-well plates as described above. After differentiation, the different ligands were added with a Biomek pin tool dispenser in serum-free DMEM overnight, next treated with 100 nM insulin. After 60 min, 1 µM puromycin was added to all wells, and the cells were incubated for an additional 30 min. The in cell western (LI-COR) was used to quantitate uptake of puromycin. Cells were fixed with 3.7% formaldehyde for 20 minutes. Non-permeabilized cell were then washed in TBS and blocked for 2 hours (Rockland MB-070) then incubated in primary anti-puromycin antibody in blocking buffer for 48 hours at 4 °C. Then the plates were washed 5 times for 5 minutes with TBS-T and then incubated with secondary antibody for 1 hr wash and analyzed using a LI-COR Odyssey imaging system.

**Protein degradation:** Rates of protein degradation were determined by measuring the release of TCA-soluble radioactivity from proteins prelabeled with [$^3$H]-phenylalanine. In brief C2C12 cells were seed in 24 well plates and after completing differentiation, myotubes were labeled with 1.0

μCi/ml of L-[3,5-$^3$H]-phenylalanine for 48 h dedifferentiation media. Cells were then treated for 24 h with ligands in DMEM containing 2 mM unlabeled phenylalanine. After treatment, the culture medium was transferred into a microcentrifuge tube containing 100 μl of bovine serum albumin (10 mg/ml) and TCA was added to a final concentration of 10% (w/v). Samples were incubated at 4° C for 1 hr followed by centrifugation for 5 min. The supernatant was used for determination of TCA-soluble radioactivity. The protein precipitates were dissolved with a tissue solubilizer (Solvable™). Cell monolayers were washed with ice-cold PBS and solubilized with 0.5 M NaOH containing 0.1% Triton X-100. Radioactivity in the cell monolayer and TCA soluble and insoluble fractions were measured using a Packard TRI-CARB 1600 TR liquid scintillation analyzer. Protein degradation was expressed as the percentage protein degraded over the 24 hr period and was calculated as 100 times the TCA-soluble radioactivity in the medium divided by the TCA-soluble plus the TCA-insoluble radioactivity in the medium plus the cell layer (i.e., myotube) radioactivity[43].

**Mitochondrial potential (Ψm):** Primary or C2C12 myotubes or mycocytes in black 96- or 384-well tissue culture plates with clear bases (Greiner Bio-One, North America, Inc.) were stained for 15 min with 200 nM MitoTracker® Orange CM-$H_2$TMRos dye (Invitrogen™ by ThermoFisher Scientific) for measuring Ψm. The cells were rinsed with steroid-free DMEM to remove un-incorporated dye, and then incubated in the differentiation media for 45 min to reach peak fluorescence. Cells were fixed in 4% formaldehyde for 20 min, stained with 300 nM DAPI for 5 min, permeabilized in PBS containing 0.1% Triton X-100 for 20 min, blocked for 1 h with 1x TBS containing 0.1% Tween-20 (TBS-T) and 2.5% normal goat serum, and incubated at 4 °C overnight with an AlexaFluor® 488-conjugated anti-skeletal muscle myosin (F59) antibody (Santa Cruz

Biotechnology, Inc. cat no. sc-32732 AF488). The next day, the myotubes were washed 4 times with TBS-T to remove unbound antibodies and rinsed twice with PBS. The stained myotubes were imaged at 10–20X magnification on the IN Cell Analyzer 6000 platform (GE Healthcare). For each treatment condition, a stack of 24–27 images containing an average of 50–100 myotubes per image were analyzed using the IN Cell Developer Toolbox image analysis software, with a customized segmentation protocol for myotubes. The average (mean) diameter and mitochondrial potential (i.e. MitoTracker staining density x area) of myotubes in these images were then calculated.

**GR nuclear translocation:** The cell line used for these experiments (3617) expresses green fluorescent protein (GFP)-tagged GR (GFP-GR) from a chromosomal locus under control of the tetracycline-repressible promoter [44]. Cytoplasm-to-nuclear translocation of the GFP-GR in response to different compounds was determined as previously described [45]. Cells were plated on a 384-well plate in DMEM medium containing 10% charcoal stripped serum (Hyclone, Logan, UT) without tetracycline (to allow the expression of the GFP-GR) at density of 2,500 cells per well. Cells were treated with compounds or a diluent as control for 30 min, fixed with paraformaldehyde, the nuclei stained with 4′,6-diamindino-2-phenylindole (DAPI) and Perkin Elmer Opera Image Screening System was used for fully automated collection of images. An image analysis pipeline was customized using the Columbus software (PerkinElmer) to automatically segment the nucleus using the DAPI channel and then construct a ring region (cytoplasm) around the nucleus mask for each cell in the digital micrographs. Translocation was calculated as a ratio of the mean GFP-GR intensity in nucleus and cytoplasm, and each value was further normalized to the value for the control (i.e., DMSO) sample on the same plate.

**Osteoblast mineralization:** Human mesenchymal stem cells (MSCs) at passage 2 were maintained in growth media consisted of alpha-MEM (Life Technologies, 32561--037) supplemented with 17% FBS (Sigma-Aldrich, 12303C), 2mM L-glutamine (Life Technologies, 25030-081), and 100 units/mL penicillin/streptomycin (Life Technologies, 15140-122). Effect of glucocorticoid receptor modulators on osteogenic differentiation of MSCs was quantified using Alizarin Red S (Sigma-Aldrich, A5533) staining. Briefly, human MSCs were seeded in the middle 8 wells of 48-well plates at plating density of 10,000 cells per well. PBS (Life Technologies, 10010-031) was added to surrounding wells to reduce evaporation of media in the middle wells during prolonged incubation. 24 h later, growth media was replaced by osteogenic induction media (OIM) consisted of low-glucose DMEM (Life Technologies, 10567-014) supplemented with 10% FBS (Sigma-Aldrich, 12303C), 50 μg/ml L-ascorbic-2-phosphate (Sigma-Aldrich, A8960), 10mM β-glycerolphosphate (Sigma-Aldrich, G9891), and glucocorticoid modulators. OIM containing test compounds was replaced every week. At the end of 2 weeks, monolayers were washed with PBS (Life Technologies, 10010-031), incubated in Richard-Allan Scientific™ Neutral Buffered Formalin (10%) (Thermo Scientific, 5725) for 2 h at room temperature, washed with deionized water and stained with Alizarin Red S for 30 min at room temperature. Monolayers were then rinsed 3 × with deionized water until clear, stain was then extracted with 10% (w/v) cetylpyridinium chloride (Sigma-Aldrich, C0732) in 10 mM sodium phosphate (Sigma, S5011 and S5136), pH 7.0 for 15 min at room temperature and the amount of extracted dye quantified spectroscopically at 562 nm. Spectroscopic analysis performed using a synergy™ NEO HTS Multi-Mode Microplate Reader (BioTek Instruments, Inc.).

**IL-6 secretion*:*** IL-1β (R&D systems) was dissolved in phosphate-buffered saline plus 0.1% bovine serum albumin (Sigma). A549 cells were plated in DMEM containing charcoal stripped serum in 384 well plates. The next day, cells were treated for 24 with GCs at 1 μM and then treated with 1 ng/ml IL-1β for 24 hrs. supernatants were collected and IL-6 assayed with AlphaLISA (PerkinElmer) using the manufacturer's instructions.

**Micro array Assay for Real-time Coregulator-Nuclear receptor Interaction (MARCoNI)**

This method has been previously described [15]. Assay mixes with 1 nM GST-tagged GR (aa 521-777, Invitrogen #PV4689), 25 nM Alexa-488 conjugated GST antibody (Fisher Scientific, #10368552), 5 mM DTT, in Coregulator buffer F (Invitrogen, #PV4547) in the presence of solvent only (2% DMSO) or 1 μM compound as indicated, were prepared on ice. Experiments were conducted on a PamStation96 (PamGene) using 2 cycles per minute at room temperature. In short, PamChip arrays (PamGene #88101) were blocked for 20 cycles with 1% Bovine serum albumin (Calbiochem #126609) in Tris-buffered saline (TBS), incubated for 40 minutes with 25 μl assay mix using 3 arrays per mix and finally rinsed twice with TBS. Binding on array (tif) images were quantified with BioNavigator software (PamGene). A modulation index, i.e. the log-transformed ratio of binding in presence over absence (solvent) of compound, was calculated for each interaction. Significance of the modulation was assessed using Student's t-Test and FDR post hoc correction.

**Antibodies**

Anti-puromycin antibody: 3RH11 (Kerafast, Inc. cat no. EQ0001), or clone 4G11 (MilliporeSigma, cat no. MABE342). Anti-AMPKα Antibody (Cell Signaling Technology, Inc.

#2532). Phospho-AMPKα (Thr172) (40H9) (Cell Signaling Technology, Inc. #2535), Anti-AKT1 (2H10) antibody (Cell Signaling Technology, Inc. #2967), Anti-pAKT-T308 (C31E5E) antibody (Cell Signaling Technology, Inc. #2965). Glut4 (3G10A3) antibody (Thermo Fisher cat no. MA5-17176). Secondary antibodies. Anti-mouse IgG goat Antibody, DyLight™ 800 (Cell Signaling Technology, cat. no. 5257). anti-Rabbit IgG (H+L) goat Antibody, DyLight 800 (Thermo Fisher Scientific cat no. SA5-10036)

**Lentiviral vectors and transduction**

Lentiviruses were produced from sets of four mouse piLenti-siRNA-GFP lentiviral vectors (Applied Biological Materials, Inc. Richmond, BC, Canada) targeting *Pelp1* (cat no. i029598), *Ncoa2* (cat no. i033105), *Ncoa1* (cat no. i038883), *Ncoa6* (cat no. i042804), *Ncor1* (cat no. i044954), or *Ncor2* (cat no. i042611), or the Control piLenti-siRNA-GFP lentiviral vector (cat no. LV015-G). HEK-293T cells in 3x 10 cm dishes per construct, were co-transfected with 10 µg lentiviral vector, 7.5 µg psPAX2, and 2.5 µg pVSV-G per dish using ProFection® kit (Promega) according to manufacturer's protocol. At ~15 h post-transfection, the media was replaced with 2.5 ml fresh media. Conditioned media containing lentiviral particles (LVM) were collected every 12 h for another 48 h, pooled, and stored at 4 °C. LVM were passed through a 0.45-micron filter to remove cell debris, and then centrifuged at 4,000 RPM overnight to pellet the viruses. The pellets were resuspended in 50 µl sterile PBS and added with 8 µg/ml polybrene to day-1 C2C12 myoblasts.

**Molecular dynamics simulations**

Initial structures used in the simulations were built from two GR LBD crystal structures, PDB

4DUC.pdb (dexamethasone-bound) and 3K23.pdb (all other states). The Modeller[46] extension within UCSF Chimera[47] was used to fill in missing parts of GR LBD in the PDB files. For apo-GR LBD, the ligand in 3K23.pdb was stripped, which was then used to dock **7** and **9** using AutoDock Vina[48]; The protonation states of residues at pH 7.4 were determined using the H++ server[49]. Ligands were parameterized using ANTECHAMBER to generate force modification files using PARMCHK2 for use in TLEAP to generate topology and coordinate files; the programs are provided in the AmberTools18 package and the steps are detailed a published tutorial (http://ambermd.org/tutorials/basic/tutorial4b). The ff14SB force field was used to describe the GR LBD. The structures were solvated in a truncated octahedral box of TIP3P water molecules with the 10 Å spacing between the protein and the boundary, neutralized with K+ and $Cl^-$ ions added to 50 mM. The system was minimized and equilibrated in nine steps at 310 K with nonbonded cutoff of 8 Å. In the first step, the heavy protein atoms were restrained by a spring constant of 5 kcal $mol^{-1}$ $Å^{-2}$ for 2000 steps, followed by 15 ps simulation under NVT conditions with shake, then two rounds of 2000 cycles of steepest descent minimization with 2 and 0.1 kcal $mol^{-1}$ $Å^{-2}$ restraints were performed. After one round without restraints, three rounds of simulations with shake were conducted for 5 ps, 10 ps, and 10 ps under NPT conditions and restraints of 1, 0.5, and 0.5 kcal $mol^{-1}$ $Å^{-2}$ on heavy atoms. Finally, an unrestrained NPT simulation was performed for 200 ps. Production runs were carried out using hydrogen mass repartitioning[50] to enable 4 fs time steps. Constant pressure replicate production runs were carried out with independent randomized starting velocities. Pressure was controlled with a Monte Carlo barostat and a pressure relaxation time (taup) of 2 ps. Temperature was kept constant at 310 K with Langevin dynamics utilizing a collision frequency (gamma_ln) of 3 $ps^{-1}$. The particle mesh ewald method was used to calculate non-bonded atom interactions with a cutoff (cut) of 8.0 Å. SHAKE was used to allow longer time steps in addition to hydrogen mass repartitioning. Production simulations were run in triplicate for 1 μs each. Analysis of trajectories was performed on the last 0.75 μs of the trajectories using CPPTRAJ and PYTRAJ[51]. For the h3-h11 distance distribution measurement, the distance (PYTRAJ dist command) between the Cα atoms of M560 on h3 and T739 on h11 was measured and plotted as a histogram. For the h3 + h11 heterogeneity measurement, the backbone Cα, C', N, and O RMSD (PYTRAJ rmsd command) of residues comprising the N-terminus of helix 3 (residues 556-565) and N-terminus of helix 11 (residues 736-743) was measured and plotted as a histogram. For the differential correlation analysis, the PYTRAJ atomiccorr command was used to generate correlation plots from the simulations of dexamethasone, **7**, and **9**, then the correlation matrices from the latter two were subtracted from the dexamethasone correlation matrix.

Dynamic network analyses were carried out after simulations by selecting Cα and C2 atoms in the protein and ligand, respectively, as nodes for networks construction. A pair of nodes was connected by their edges if they have satisfied a distance requirement (<4.5 Å) for at least 75% of the simulation time. Carma program and NetworkView plugin in VMD were utilized for producing the dynamic network by calculating the cartesian covariance and correlation between two nodes[52,53]. The edge distances were derived from pairwise correlations as a measure of communication within the network. Suboptimal paths between the C-terminus of h5 and h12 were identified using the Floyd-Warshall algorithm and analyzed by subopt program in the VMD NetworkView plugin.

**Data Availability Statement**

The raw and processed nascent RNA-seq dataset generated in this study has been deposited in the National Center for Biotechnology Information (NCBI) Gene Expression Omnibus (GEO), with the accession code GSE149453. Other mRNAseq data cited in this work was downloaded from the GEO with accession code GSE124636[54]. Sample *R* scripts used are available at https://github.com/jnwachuk/ML_in_GR_signaling_networks.
