## Supplementary material for "Chemical Systems Biology Reveals Mechanisms of Glucocorticoid Receptor Signaling": Methods

Supplementary Tables: Pages 2–4
Supplementary Figures: pages 5–6
Supplementary Notes: page 7
Supplementary References: pages 8–10

### Supplementary Table

| Gene | Function / Rationale | Refs |
|---|---|---|
| *Acss1* | important for maintaining body temperature during fasting, energy homeostasis, and energy expenditure under ketogenic conditions; converts acetate to acetyl-CoA in the TCA cycle | 55 |
| *Bcl2l1* | encodes proteins located in the outer mitochondrial membrane; regulates VDAC opening to control mitochondrial potential, ROS production, and cytochrome C release | 56,57 |
| *Ctgf* | has important roles in cell adhesion, migration, proliferation, angiogenesis, skeletal development, and tissue wound repair; is critically involved in fibrotic disease | 58 |
| *Ddit4* | negative regulator of mTOR; associated with nutrient deprivation in skeletal muscle. | 59,60 |
| *Deptor* | inhibits mTORC1 and mTORC2 activity | 61 |
| *Eif4a1* <br> *Eif4a2* | RNA helicase subunits of the eIF4F complex involved in mRNA cap recognition and binding to ribosome; involved in mTOR and TGF-β signaling pathways, and skeletal muscle atrophy | 62 |
| *Errfi1* | Negative regulator of the EGFR family; inhibits EGFR activity; modulates steroids response | 63 |
| *Fbxo32* | also known as MAFbx or Atrogin-1; a muscle-specific gene required for muscle atrophy | 64 |
| *Figf* | c-fos-induced growth factor/vascular endothelial growth factor D (Figf/Vegf-D) | 65 |
| *Fkbp5* | regulates AKT2 activity and metabolic function. | 25 |
| *Foxo3* | causes skeletal muscle atrophy by inducing the Atrophy-Related Ubiquitin Ligase, Atrogin-1 | 66 |
| *Irs1* | coordinates skeletal muscle growth and metabolism via the AKT and AMPK pathways. | 67 |
| *Klf15* | is upregulated by fasting and decreased by feeding and insulin via PI3K signaling | 68 |
| *Map3k8* | Activates IκB kinases; promotes nuclear localization of NF-κB; promotes muscle atrophy | 69,70 |
| *Nfkbia* | reduces NF-κB activity that can contribute to insulin resistance and muscle atrophy | 71 |
| *Nr4a3* | negative control | |
| *Pdk4* | Plays a pivotal role in control of metabolic flexibility in skeletal muscle | 72 |
| *Pld1* | regulates muscle cell size via the activation of mTOR signaling | 73 |
| *Ppp2r2c* | Ser/Thr phosphatase; competes against mTOR for IRS-1; regulates IRS-1 phosphorylation | 74 |
| *Prkar2b* | PKA subunit; disruption of PKA in mice promotes healthy aging | 75 |
| *Pax7* | Pax-7 upregulation inhibits myogenesis and cell cycle progression in satellite cells | 76 |
| *Rhoj* | RhoJ is an endothelial cell-restricted Rho GTPase that mediates vascular morphogenesis and is regulated by the transcription factor ERG; orchestrates cell-cell communication | 77,78 |
| *Rpsa* | negative control | |
| *Sgk1* | regulates muscle mass maintenance by suppressing proteolysis and autophagy, and increasing protein synthesis; important to maintain skeletal muscle homeostasis and prevent atrophy | 79 |
| *Socs2* | negatively regulates growth hormone action in vitro and in vivo; inhibits mitochondria biogenesis via inhibiting p38 MAPK/ATF2 pathway in C2C12 cells | 80,81 |
| *Syk* | May be catabolic through inhibition of PKA and Creb1 activity | 42, 82 |
| *Trim63* | Muscle-specific E3 ubiquitin ligase involved in muscle atrophy and cancer cachexia | 83 |
| *Tsc22d3* | Regulates MyoD/HDAC1 transcriptional activity; mediates the anti-myogenic effect of GCs | 84 |
| *Vav2* | Rho GTPase; modulates muscle mass and glucose uptake into skeletal muscle | 85 |
| *Vegfc* | Overexpression of VEGF-C induces weight gain and insulin resistance in mice | 86 |

### Supplementary Table 1. Genes examined using nanoString nCounter

| **Phenotype** | Glut4 | Degradation | Ψm | Synthesis | pAKT |
|---|---|---|---|---|---|
| [#]*Predictive capacity* | 84% | 46% | 68% | 70% | 52% |
| **Predictors** | **Fkbp5* | *Ψm | *Degradation | *CHD9_1023 | *Synthesis |
| | **Socs2* | **Socs2* | *Synthesis | *NCOR2_2330 | *PRAME_91 |
| | **Bcl2l1* | Synthesis | CNOT1_557 | **Socs2* | *NCOR2_2123 |
| | **Tsc22d3* | PRAME_91 | CHD9_1023 | *PRAME_91 | |
| | *NCOA1_677 | | NCOR2_2330 | **Fbxo32* | |
| | *NCOA1_737 | | *Socs2* | **Sgk1* | |
| | *Sgk1* | | | **Irs1* | |
| | NRBF2_128 | | | NCOR1_2251 | |
| | PPARGC1B_338 | | | Ψm | |
| | *Vav2* | | | NELFB_328 | |
| | NCOA2_733 | | | pAKT | |
| | PELP1_446 | | | CHD9_855 | |

**Supplementary Table 2. Significant predictors of GR-mediated phenotypes**

Compounds **1-3**, **6-27** were profiled for regulation of specific target genes in skeletal muscle, and GR interaction with specific peptides. Significant predictors were identified using the random forest classifier[21] and statistical thresholds established with the Boruta wrapper algorithm for feature selection[22]. The number in a peptide name indicates the N-terminal amino acid of the peptide in the full-length protein. *Minimal set of predictors; [#]The predictive capacity of the models, is the fraction of variance in the dependent variable that is explained by the minimal predictive model.

| Peptide | Synthesis | Ψm | Degradation | pAKT | Glut4 | *Bcl2l1* | *Fkbp5* | *Sgk1* | *Tsc22d3* | *Vav2* |
|---|---|---|---|---|---|---|---|---|---|---|
| PELP1_446 | 0.22 | 0.02 | 0.10 | 0.45 | 0.65 | 0.89 | 0.96 | 0.86 | 0.96 | 0.87 |
| NCOA1_737 | 0.24 | 0.02 | 0.12 | 0.46 | 0.69 | 0.88 | 0.92 | 0.78 | 0.97 | 0.89 |
| NCOA1_677 | 0.21 | 0.01 | 0.10 | 0.43 | 0.71 | 0.87 | 0.91 | 0.74 | 0.96 | 0.92 |
| NCOA2_733 | 0.26 | 0.02 | 0.13 | 0.50 | 0.73 | 0.85 | 0.90 | 0.72 | 0.97 | 0.92 |
| NRBF2_128 | 0.18 | 0.00 | 0.10 | 0.38 | 0.65 | 0.83 | 0.89 | 0.70 | 0.93 | 0.89 |
| PPARGC1B _338 | 0.21 | 0.01 | 0.13 | 0.47 | 0.75 | 0.83 | 0.82 | 0.66 | 0.93 | 0.88 |
| NCOR2_2230 | 0.62 | 0.41 | 0.21 | 0.34 | 0.02 | 0.03 | 0.37 | 0.12 | 0.12 | 0.05 |

**Supplementary Table 3. GR interactions with coregulator peptides selectively predict GR-mediated gene expression and skeletal muscle phenotypes.**

Linear regression analysis comparing the predictive power of the indicated peptide interactions for the indicated skeletal muscle phenotypes and target genes. Coefficient of determination, $r^2$, are shown.

### Supplementary Figures

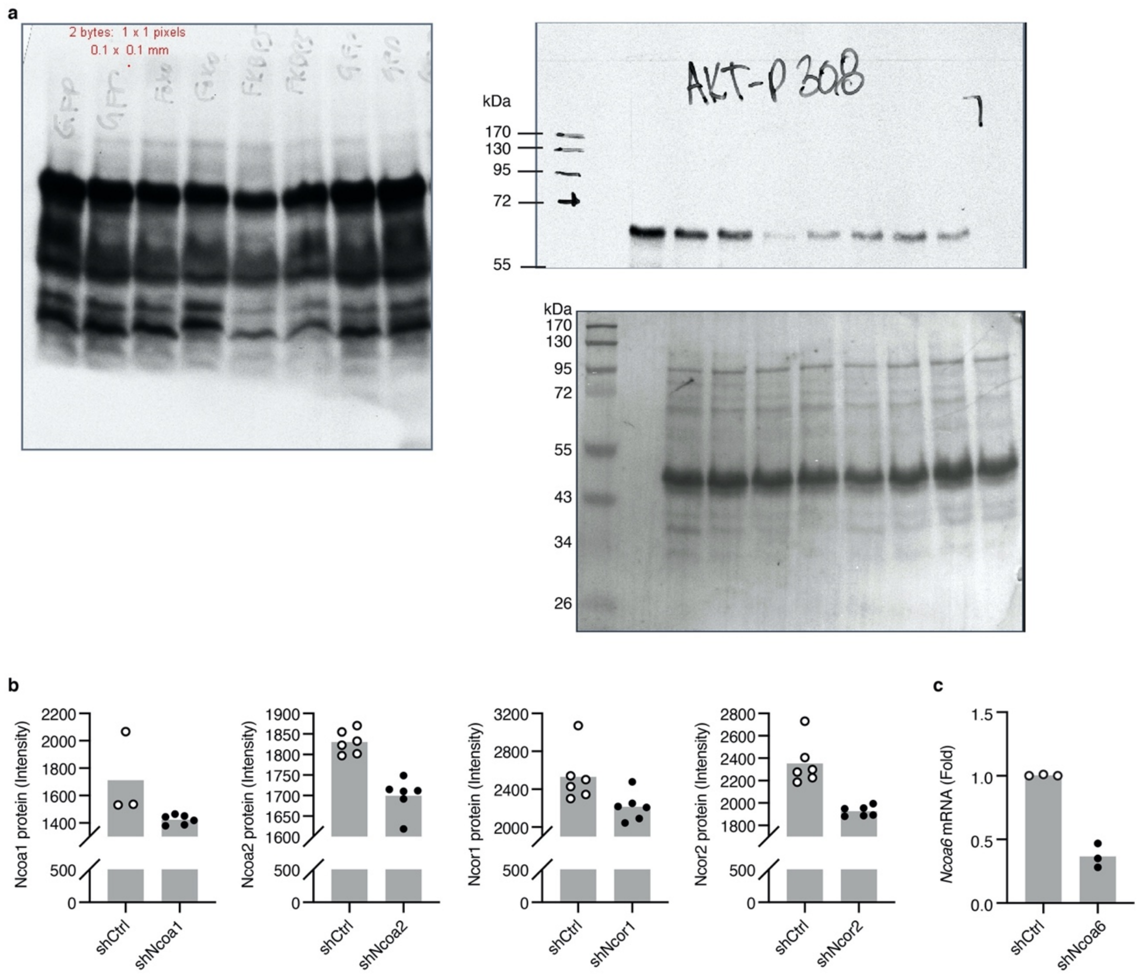


**Supplementary Figure 1. Additional information from gene perturbation studies**

**a**) Full scanned images of Western blot films presented. See **Figure 3b-c**.

**b-c**) shRNA knockdown controls. Lentiviral transduction with control or the indicated shRNAs were used to generate stable C2C12 cell lines. **b**) Effects of the shRNA knockdown on the indicated proteins were compared by in cell western (LICOR Odyssey). Bars represent the mean; n=6, except for Ncoa1/shCtrl where n=3 biologically independent samples. **c**) Knockdown of Ncoa6 mRNA was verified in a shNcoa6 C2C12 stable cell line by qPCR. Bars represent the mean; n=3 biologically independent samples.

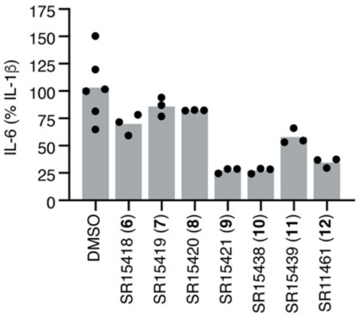


**Supplementary Figure 2. Effects of C3-substituted GCs on IL-6 secretion by A549 cells.** Bars represent the mean; n=3, except for DMSO where n=6 biologically independent samples. See also **Extended Data Fig. 4**

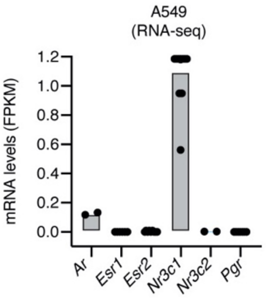


**Supplementary Figure 3. Expression of steroid receptor mRNAs in a published A549 cell RNA-seq dataset**[54]. Only *Ar* which encodes AR, and *Nr3c1* which encodes GR, are expressed in these cells.
